# Neuronal Hyperexcitability and Seizure Susceptibility in a Novel *Kcnt1 G269S* Mouse Model

**DOI:** 10.1101/2025.07.22.666003

**Authors:** Sarah L. Olguin, Ammara Rehman, Daniel Davis, Sean W. Berg, Elizabeth Sahagun, Payton S. Oswalt, Mandar Patil, Paula D.M. Sullivan, Olivia Y. Haouchine, Luther Ortega, Bing Zhang, Jill L. Silverman, Elizabeth C. Bryda, Roy Ben-Shalom

## Abstract

*KCNT1*-related epilepsy is an autosomal dominant neurodevelopmental disorder resulting from *de novo* pathogenic variants in *KCNT1* which encodes a sodium-gated potassium channel. To date, 64 *KCNT1* mutations have been identified across 248 individuals. The majority of pathogenic *KCNT1* mutations are gain-of-function (GOF), with each exhibiting unique channel biophysical properties, channel dynamics, electrophysiological spiking, and epileptiform characteristics. These diverse neuronal electrical profiles, contribute to a spectrum of epilepsy syndromes ranging from severe early-onset epileptic encephalopathies to focal epilepsies with milder outcomes. Among these, variants at Gly288 are recurrent, reported in 29 of 248 individuals in the largest published cohort to date, and 79% (23/29) of whom presented with severe early-onset epilepsy, yet no *in vivo* model of this pore-domain variant currently exists.

Our multi-disciplinary collaboration generated a novel mouse model (C57BL/6-*Kcnt1^em1Bryd^*) that carries the G269S variant of *Kcnt1*, corresponding to the human G288S variant, located within the coding region of the channel pore. We characterized network excitability of primary cultured cortical neurons from all three genotypes (*Kcnt1*^+/+^, *Kcnt1*^+/G269S^, and *Kcnt1*^G269S/G269S^) on high-density microelectrode arrays, since many *KCNT1* mouse models require homozygosity for phenotypic detection.

*Kcnt1*^+/G269S^ neurons exhibited sustained hyperexcitability and hypersynchronous bursting versus *Kcnt1*^+/+^ controls, while *Kcnt1*^G269S/G269S^ neurons showed early excessive bursting followed by a progressive decline in coordinated network activity by 28 days-in-vitro. Therefore, behavioral comparisons were conducted between *Kcnt1*^+/G269S^ and *Kcnt1*^+/+^ mice including developmental milestones, motor and cognitive abilities, respiratory physiology, and seizure susceptibility. *Kcnt1*^+/G269S^ mice exhibited spontaneous behavioral seizures, poor motor coordination, erratic breathing, and increased apneas, as functional outcome measures. Critically, *Kcnt1*^+/G269S^ mice were more susceptible to thermal-induced seizures in early life and chemoconvulsant induced seizures in adulthood.

In summary, these data: (i) generated and validated a novel mouse model of *KCNT1*-related epilepsy, (ii) provide strong *in vitro* evidence of neuronal hyperexcitability, (iii) illustrate behavioral spontaneous seizures, early-life seizures, and adult seizure susceptibility as functional outcome measures, (iv) elucidated alterations in exploratory locomotion and poor motor coordination, (v) identified increased erratic breathing in female and increased apneas in male *Kcnt1*^+/G269S^ mice and (vi) lay the groundwork for future analysis of neural activity *in vivo* and modeling circuit level dynamics *in vitro*, *in vivo,* and *in silico*.

## Introduction

Genetic studies have shown that approximately one-third of neurodevelopmental disorders (NDDs) are associated with gene mutations affecting neuronal ion channels, emphasizing their critical role in brain pathology.^1–3^ To understand how these mutations alter channel kinetics, *in vivo* experimental models are essential.^4–7^ The *KCNT1* gene encodes a sodium activated potassium (K_Na⁺_) channel that regulates neuronal excitability by influencing the resting membrane potential and hyperpolarization following action potentials. Mutations in *KCNT1* result in diverse electrophysiological profiles, that contribute to a spectrum of epileptic syndromes ranging from severe early-onset epileptic encephalopathies such as Epilepsy of Infancy with Migrating Focal Seizures (EIMFS), to focal epilepsies such as Autosomal Dominant Nocturnal Frontal Lobe Epilepsy (ADNFLE), West syndrome, and Ohtahara syndrome.^8^ Notably, the same variant can present across multiple syndromes, and no strict genotype-phenotype correlation has been established.

To date, 64 *KCNT1* mutations have been identified across 248 individuals,^9^ including our mutation of interest p.Gly288Ser.^10–12^ While, most of the mutations identified are located within the intracellular C-terminal region, missense mutations at p.Gly288Ser are located within the coding region of the channel pore.^11,12^ Variants at Gly288 are among the most recurrent in the disorder, reported in 29 of 248 individuals in the largest published cohort to date, 23 of whom (79%) presented with severe early-onset epilepsy (EIMFS or non-EIMFS developmental and epileptic encephalopathy) underscoring the clinical importance of this position.^9^ p.Gly288Ser was first identified in two unrelated patients with malignant migrating partial seizures in infancy (MMPSI) characterized by seizures beginning before 6 months of age, migrating focal motor seizures, nearly continuous multifocal seizures migrating between cortical regions and hemispheres, delayed psychomotor development, and, critically, resistant to antiepileptic drugs.^13^ An additional individual with MMPSI and a p.Gly288Ser mutation had onset of seizures at 2 months of age that were characterized as motor seizures during sleep.^14^ Furthermore, individuals with missense mutations at p.Gly288Ser have also been diagnosed with temporal lobe epilepsy, epilepsy with tonic-clonic seizures, multifocal epilepsy, and epilepsy with myoclonic-atonic seizure phenotype, further demonstrating the unmet need of a validated *in vivo* model of G288S pathogenic variant.^9,10,15,16^

The gain-of-function nature of the G288S variant has been established across multiple model systems. D*rosophila melanogaster* lack a *KCNT1* ortholog, allowing for the introduction of the human *KCNT1* (*hKCNT1)* gene. While the wildtype *hKCNT1* gene showed no phenotype, expression of gain-of-function (GOF) human variants exhibited seizure activity rates of 41% (G288S), 48% (R398Q), and 38% (R928C), respectively.^17^ Zhang’s laboratory illustrates that *hKCNT1* G288S and R928C mutations increased potassium conductance, impaired larval locomotion, and hyperpolarized the resting membrane potential.^18,19^ Consistent with these findings, expression of the G288S variant in HEK cells exhibited: (i) a 21-fold increase in K_Na⁺_ current amplitude; (ii) an approximate 20% increase in channel open probability; (iii) a hyperpolarizing shift in activation threshold, reducing voltage sensitivity; and (iv) minimal changes in channel kinetics.^10^ Despite this convergent evidence that G288S is a strong GOF variant, an appropriate *in vivo* model has been lacking. While mouse models of other *Kcnt1* mutations exist, outlined in Supplemental Table 1, most require homozygosity for phenotypic detection. Importantly, each variant effects channel kinetics by differentially altering the activation threshold, open probability and current amplitude^10^ impacting neuronal excitability and seizure susceptibility dependent on the variant, neuronal subtype, and background strain of the rodent model^20–22^ adding complexity to understanding circuit-level dysfunction in *KCNT1*-related epilepsies. Critically, affected individuals are heterozygous, yet no heterozygous model of a pore-domain variant has been characterized. These data support the generation of the *Kcnt1* GOF G288S (G269S in mice) experimental model, termed B6-*Kcnt1^em1Bryd^*. This model enabled targeted characterization of electrophysiological properties *in vitro* and functional outcomes *in vivo* providing a valuable tool for developing therapeutics for *KCNT1*-related epilepsy.

## Materials and methods

### Generation of the G288S Variant (G269S Mouse Model)

All experimental procedures were performed in accordance with the National Institutes of Health (NIH) Guide for the Care and Use of Laboratory Animals and approved by the Institutional Animal Care and Use Committees (IACUC) at the University of Missouri and University of California, Davis School of Medicine. The novel *Kcnt1* mutant mouse model was generated by the MU Animal Modeling Core (Columbia, MO) using a standard knock in method involving CRISPR/Cas9 genome editing. The strategy for generating the *Kcnt1* exon 11 codon change was to replace the wild-type (WT) mouse *Kcnt1* gene codon GGG with AGC creating a missense mutation. Glycine (G) is replaced by serine (S) at amino acid position 269 in the mouse KCNT1 protein which mimics the G288S human variant. C57BL/6J (stock #000664) animals were purchased from the Jackson Laboratory (Bar Harbor, ME). Super ovulated females (4-5 weeks of age) were mated to stud males (∼10 weeks of age) overnight and zygotes were collected the following morning. sgRNAs were ordered as chemically modified synthetic sgRNAs through Synthego (Redwood City, CA). The chemical modifications on each sgRNA were 2′-O-methyl analogs and 3′ phosphonothioate internucleotide linkages at the first three 5′ and 3′ terminal RNA residues. The repair template was chemically synthesized by IDT (Coralville, IA) as a 200bp single-stranded DNA oligo (ssODN). The ssODN was complementary to the non-target strand and contained symmetrical homology arms. An additional silent mutation was introduced for genotyping purposes.

#### sgRNA sequence

5′-TCCAGGTGCTGGATCCCAC-3′. Repair template sequence (sgRNA sequence disrupted by desired mutation): 5′-CTGCTGCAGCCTCTCCCCCACTGCTGCAGCCCCAGCCCCACTGCTGCAGCCCCAGCC CCACTGCTGCAGCCCTGATCTTCCTGTTTCTAGGACCTGT**A**G**C**ATCCAGCACCTGGA GCGAGCAGGTGGCAACTTGAACCTGCTGACCTCCTTCTACTTCTGCATCGTGACTTTC TCAACTGTGGGCTTCGGTGATGTGAC -3′. A mix containing a final concentration of 3.0μM sgRNA, 2.0μM enhanced specificity Cas9 protein (Sigma) and 1.6μM ssODN was made immediately prior to electroporation. CRISPR sgRNA/Cas9 RNP complexes were formed by gently mixing the sgRNA and Cas9 protein together and incubating at room temperature for 10min. After RNP formation, the ssODN repair template was added to the mixture. Using a NepaGene21 electroporator along with a 1.5-mm gap glass slide electrode, zygotes were electroporated under the following conditions: Poring pulse: 40V, 3.5ms length, 50ms interval, 10% decay rate, positive polarity (x4 pulses) Transfer pulse: 5V, 50ms length, 50ms interval, 40% decay rate, alternating polarity (x5 pulses). Manipulated zygotes were then surgically transferred to pseudo-pregnant surrogate CD1 females (8 weeks of age) purchased from Charles River Laboratories (Wilmington, MA). Multiple animals carrying the desired single base pair change were identified by genotype analysis and confirmed by nucleotide sequence analysis of the region around the base pair change. The founders confirmed to carry the mutation were backcrossed to C57BL6/J for 2 generations and then to C57BL6/NJ (Jackson laboratory #005304) for 2 generations to establish the strain C57BL/6NJ *Kcnt1^em1Bryd^* which is available at the Mutant Mouse Resource and Research Center at the University of Missouri (MMRRC#075727) and is available upon request at www.mmrrc.org. In 2021, animals were transferred to UC Davis, School of Medicine (Sacramento, CA) to establish a colony. This colony was maintained by backcrossing to C57BL/6J for >10 generations.

#### Genotyping

Genomic DNA was extracted by placing a small amount of tissue (ear or tail) in 100µl alkaline lysis reagent (25mM NaOH and 0.2mM EDTA at a pH of 12) and heating at 95 °C for 1hr. This was followed by addition of 100µl neutralization buffer (40mM Tris-HCl) with vortexing for 5secs to mix. The <u>PCR primers were Kcnt1.F1</u>: 5’-cattccacactgcagccctgtctct-3’and <u>Kcnt1.R1</u>: 5’-caagggtgacacagatcaggatgaccac-3’. The 25µl reaction contained the following: <u>12.5µl</u> EmeraldAmp^®^ Supermix (Takara Bio USA Inc., San Jose, CA); <u>10.0µl</u> H_2_O; <u>0.75µl</u> Forward primer (10µM); <u>0.75µl</u> Reverse primer (10µM); <u>1µl</u> genomic DNA (∼5-50ng). Thermal cycler conditions were 1) 95°C, 3mins, 2) 95°C, 30secs, 3) 69°C, 30secs, 4) 72°C, 1min. Steps 2-4 were repeated 40 times, 5) 72°C, 7mins, 6) 4°C, hold. The expected PCR amplicon = 403bp. Following PCR amplification, 2.8µl CutSmart buffer and 1.0µl *BtsC*I (20 units; New England Biolabs, Ipswich, MA) were added directly to the PCR reaction and incubated at 50°C for 2hrs. Gel electrophoresis was performed with a 3% agarose gel in 1X TAE buffer at 80V for 1hr. Expected restriction fragment sizes: WT allele = 19bp, 23bp, 42bp, and 319bp; mutant allele = 19bp, 23bp, 42bp, 113bp, 206bp. The 319bp fragment in the WT allele is cut into 113bp and 206bp fragments in the mutant allele.

### Primary Neuronal Culture

Test mice were generated by crossing male and female *Kcnt1^+/G269S^*to yield pups of three genotypes (*Kcnt1^+/+^, Kcnt1^+/G269S^*, and *Kcnt1^G269S/G269S^*) and neonatal cortices (PND 0-2) were collected. Primary cortical neuron-glia cultures were prepared as previously described.^23,24^ Briefly, cortices were minced with a razor blade prior to incubation in Hibernate A containing papain and DNase. The tissue was then triturated with glass pipettes to dissociate cells from the extracellular matrix (Bellco Glass, Vineland, NJ). Dissociated cells were counted and plated as described below.

#### High Density Microelectrode Array Plating

MaxWell Biosystems high density microelectrode arrays (HD-MEAs) were used to assess electrophysiological activity of primary neuronal cultures, as previously described.^25,26^ MaxTwo 6-well HD-MEA plates were sterilized, filled with 2mL of plating medium (Neurobasal Plus medium (Gibco, ThermoFisher Scientific), 2% B27 Plus supplement (×50) (Gibco, ThermoFisher Scientific), 1x GlutaMAX supplement (Invitrogen), 10% heat-inactivated horse serum (Invitrogen), and HEPES buffer (1M, pH 7.5) (Fisher Scientific)), then incubated for 2 days at 37°C with 5% CO_2_. On the day of cell plating, the plating media was removed, and chips washed with sterile water, coated with primary coating (50μl of 0.07% polyethyleneimine (PEI, Sigma-Aldrich)), and incubated for 1hr. The primary coating was removed, the chips were washed 3 times with sterile water, and vacuum dried. Secondary coating (50ul Laminin (0.02 mg/mL; Sigma-Aldrich)) was applied and incubated (∼ 1hr) until ready to plate cells. The laminin coating was removed, 50uL of primary cortical cells were added to the HD-MEAs, yielding approximately 125,000 cells per chip, and incubated for 1hr before the addition of 2mL plating medium. 24 hours later, 50% of media was removed and replaced with cell culture maintenance media (Dulbecco’s Modified Eagle Medium (DMEM) supplemented with GlutaMax, D-glucose and sodium pyruvate (Gibco, ThermoFisher Scientific) and 10% horse serum). Media changes occurred twice weekly by removing 50% of the media and replacing it with an equal volume of cell culture media. Neural activity was recorded twice a week as described below.

#### HD-MEA Electrophysiological Recordings and Data Processing

HD-MEA recordings were performed between days *in vitro* (DIV) 3 to 30 to capture the developmental trajectories of dissociated neuronal cultures. Due to differences in recording schedules across experiments, some DIVs varied slightly between litters (e.g., DIV13 vs 14, DIV27 vs. 28). Each session began with the Activity Scan Assay, which recorded 30sec segments across the entire chip under ‘Sparse 7x’ configuration to assess overall activity of the chip. Based on spike amplitude and spatial distribution, 1,020 electrodes were selected from regions exhibiting high activity (spike amplitude) to maximize neuron coverage in the Network Scan Assay. Spontaneous activity was recorded for 300sec at a 10kHz sampling rate using the MaxTwo device (MaxWell Biosystems).

Data processing focused on two aspects: (1) The activity of the culture over time, and (2) network-level dynamics reflecting population synchrony and communication. Raw extracellular signals were bandpass filtered between 300Hz and 3,000Hz to isolate multi-unit activity (MUA). Spike detection was performed using a threshold set at 5X the standard deviation of baseline noise. A recording channel was considered active if it exceeded a minimum firing rate of 0.1Hz and spike amplitude threshold of 20µV. The mean firing rate and mean spike amplitude were computed from all active channels to assess the overall culture health and neuronal proliferation. To characterize network-level activity, detected spikes were transformed into spike raster plots and binned to generate population firing rates. A Gaussian kernel was applied to these binned spike trains to estimate a continuous measure of population activity, capturing burst dynamics and synchronous events. Statistical comparisons between genotypes were conducted using one-way ANOVAs (pooled DIV13 to 21 recordings for Fig. 2; individual DIVs for Fig. 3) and a two-way ANOVA (genotype × DIV) across development for Fig. 3, each followed by Tukey’s post hoc tests; N per genotype, F values and exact post hoc p values are listed in the figure legends and in Supplemental Table 3. To ensure analytical consistency, adjacent DIVs with ±1 day variation were grouped during analysis. All analyses were conducted using custom MATLAB R2024 scripts in combination with the MaxWell Biosystems Toolbox (v23.2).

For the pharmacological modulation experiment, HD-MEA recordings were performed at DIV23 in *Kcnt1*^+/+^ and *Kcnt1*^+/G269S^ cortical cultures. Wells were assigned to one of two groups, Vehicle Control or Treatment. Each recording session began with a full Activity Scan Assay to assess overall chip activity, followed by a 600 sec Network Scan Assay at a 10 kHz sampling rate using the MaxTwo device (MaxWell Biosystems), establishing the Baseline recording. Gabazine (10 µM), a GABA-A receptor antagonist, was then added to Treatment wells, and an equivalent volume of water was added to Vehicle Control wells; 1 hr later, a second 600 sec network recording was performed. AP5 (20 µM) and NBQX (20 µM), which block NMDA and AMPA/kainate receptors, respectively, were then added to the same Treatment wells, and a second equivalent volume of water was added to Vehicle Control wells; 1 hr later, a third and final 600 sec network recording was performed. This yielded three sequential 600 sec recordings per well (Baseline, Gabazine 1 hr, AP5+NBQX 1 hr), with n = 3 wells per genotype per group. Network burst detection, participation fraction, and peak population firing rate were computed as described above. Statistical comparisons between groups were conducted using a mixed-design ANOVA (group × condition, with group as a between-subjects factor and condition as a within-subjects factor) followed by post hoc unpaired t-tests comparing each well’s change from Baseline between Treatment and Vehicle Control groups.

### Subjects for Developmental Milestones, Behavioral Assays, and Seizure Susceptibility

All mice were housed in a temperature- and humidity-controlled vivarium maintained on a 12:12 light-dark cycle and tested during the light cycle. Animals were fed a standard diet of Teklad global 18% protein rodent diets 2918 (Envigo, Hayward, CA, USA). Pups were labeled by paw tattoo over postnatal days (PND) 2-4 using non-toxic animal tattoo ink (Ketchum Manufacturing Inc., Brockville, ON, Canada) and tail biopsies (1–2 mm) were taken for genotyping, following the UC Davis IACUC policy regarding tissue collection (PND 4-7). Mice were weaned at PND 21, and group housed by sex with 2-4 animals per cage. Assays were performed in order of least to most stressful and between the hours of 9:00AM PST and 4:00PM PST during the light phase. All behavior testing was conducted by an experimenter blinded to genotype and included both sexes, animals were randomized by cage for testing order each day. Mice were allowed to habituate in their home cages in a dimly lit (50lux) room adjacent to the testing room for 1hr prior to the start of testing to limit any effect of transporting between the vivarium and testing suite. Between subjects, testing apparatus surfaces were cleaned using 70% ethanol and allowed to dry. Six Cohorts of mice were used for a combination of developmental milestones, seizure-related outcomes, behavioral phenotypes, and respiratory physiology, with full details below.

### Behavioral Assays

Cohort 1 comprised 7 litters (n=44 mice; 8 female and 14 male per genotype). A subset (5 litters; 5 female and 6 male Kcnt1+/+; 6 female and 8 male Kcnt1+/G269S) was assessed for developmental milestones on PND 3, 5, 9 and 11 (Supplemental Table 2). The full cohort was then tested in this order: open field, rotarod, novel object recognition, contextual and cued fear conditioning. See **Supplemental** for developmental milestones, novel object recognition, and fear condition methods. Cohort 2 comprised 4 litters (n=33 mice; 5 female and 7 male *Kcnt1^+/+^;* 10 female and 11 male *Kcnt1^+/G269S^*) and Cohort 3 comprised 5 litters (n=28 mice; 8 female and 4 male *Kcnt1^+/+^;* 9 female and 7 male *Kcnt1^+/G269S^* a replicate Cohort of Cohort 2) were tested during a critical time window (PND 21-23) for hyperthermia-induced seizures. During this same time window, spontaneous seizures were observed within an independent cohort, Cohort 4, comprised of 12 litters (n=56 mice; 13 female and 10 male *Kcnt1^+/+^;* 18 female and 15 male *Kcnt1^+/G269S^*). Adult seizure assessment occurred by high dose (lethal) chemoconvulsant seizure induction with pentylenetetrazol (PTZ) intraperitoneally (i.p.) in behaviorally naïve Cohort 5 mice comprised of 10 litters (n=22 mice; 5 female and 5 male *Kcnt1^+/+^;* 9 female and 3 male *Kcnt1^+/G269S^*). Finally, to further investigate *Kcnt1*’s potential influence on respiratory physiology, Cohort 6 mice comprised of littermates from 8 litters (n=42 mice; 11 female and 9 male *Kcnt1^+/+^;* 9 female and 13 male *Kcnt1^+/G269S^*) that underwent whole body plethysmography (WBP).

#### Open Field Locomotion

Exploratory locomotion in a novel open field environment was assayed in a room illuminated at ∼40lux in a 40cm×40cm×30.5cm arena as previously described.^27–30^ Exploratory locomotion is an essential assay for gross motor abilities and physical activity over time, identifying key conserved neuronal circuit and motoric phenotypes from flies to humans.

#### Rotarod

Motor coordination, balance, and motor learning were assessed using an Ugo-Basile accelerating rotarod (Stoelting Co, Wood Dale, IL) as previously published.^31,32^ Mice are placed on a rotating cylinder that slowly increases from 4 to 40 revolutions per min (rpm) over a 5min trial, with 3 trials per day using a 60min inter-trial interval, and subsequently repeated over 3 consecutive days for a total of 9 trials. Performance was scored as latency to fall (sec) from the cylinder with a maximum latency of 300sec.

#### Whole Body Plethysmography (WBP)

Whole-body plethysmography (WBP) chambers (Buxco/FinePointe, Data Sciences International, St. Paul, MN) were utilized to assess respiratory physiology in Cohort 6 subjects. Breath-by-breath parameters were quantified as well as the occurrence and time spent in apneas (an absence of inhaling or exhaling). A normal breath is a breath that exhibits an identifiable inspiratory and expiratory phase. Type 0 apneas are an extended pause in breathing that were not immediately preceded by an augmented breath (a distinct increase in breath volume relative to the immediately preceding breaths). An erratic breath is a breath that exhibits an unequal inspiratory volume and expiratory volume, is not an apnea, nor an augmented breath. Given that age-matched males are larger than females in body weight and size, both contributing parameters that influence the volume of air respirated during the plethysmograph diagnosis subjects were sex-separated to eliminate sex-originating variables for WBP presentation and analysis.

#### Hyperthermia-induced Seizure Susceptibility

Postnatal day (PND) 21-23 subjects were utilized to reflect the same approximate vulnerable developmental window as the human population.^33,34^ Subjects were habituated in an adjacent room for at least 30mins in low light (∼30 lux). Subjects were given a brief anesthetic exposure (0.1ml 4% isoflurane for 30sec-1min until loss of consciousness) for insertion of rectal temperature probe (Physitemp RET-4, Clifton, NJ, USA), prior to placement in the testing arena. A heat lamp placed above the arena was positioned to maintain an internal body temperature of 37.5°C for a 10min habituation period. Following arena habituation, the distance of the heat lamp was dynamically adjusted to avoid rapid overheating and gradually raise the internal body temperature evenly by approximately 0.5°C every 2mins. In order to promote survivability post-heat exposure testing continues until a behavioral seizure score of 3 or above is observed or until the body’s internal temperature reaches 42.5°C, in which case the heat lamp is immediately turned off and distanced from the arena to allow the subject to cool to 40°C proceeded by removal. An observer blind to genotype records the type of seizure using a modified Racine Scale (1. Staring, 2. Head nodding, 3. Unilateral forelimb clonus, 4. Bilateral forelimb clonus, 5. Rearing and falling, 6. Clonic seizure) previously described.^35^ The febrile Racine Scale is unique in that animals typically progress through the stages over time and then recover once they are removed from the heat source. Animals are returned to their home cage and provided hydrogel once they are active and exhibit normal ambulatory behavior (∼30mins).

#### Spontaneous Seizure Observations

Seizures were observed following temporary intra-facility room transfers, from the vivarium to the behavioral suites. A spontaneous seizure event was defined as erratic motor activity, loss of righting, generalized tonic-clonic seizure, and death. Handling precautions were maximized to reduce any additional cage disturbances/stress.

### Adult Seizure Susceptibility by a Lethal Dose of the Chemoconvulsant Pentylenetetrazol (PTZ)

Dosing was conducted between 9am-2pm in a well-lit room and followed our validated dosing regimens for C57BL/6J. Subjects were weighed and administered PTZ (70mg/kg, i.p.), then placed in a clean, empty cage with subsequent seizure stages live scored using a modified Racine scale for a maximum of 30mins. Stages scored were: (1) first jerk/Straub’s tail, (2) loss of righting reflex, (3) generalized clonic-tonic seizure, (4) full tonic extension, (5) death. The PTZ Racine Scale is unique in that animals typically progress through the stages over time.

### Statistical Analysis

Data were analyzed with GraphPad Prism by two-tailed Student’s t-test for single parameter comparison between the two genotypes or a Two-Way Repeated Measures ANOVA for comparison of genotype over time (e.g. development, open field, rotarod, pre-and post-fear conditioning). Significant ANOVAs were followed by Tukey’s post hoc testing for network measures and by Bonferroni’s post hoc testing for the adult PTZ seizure stage progression and NOR. Categorical outcomes, including probability of survival, were compared by chi-square test and survival was analyzed by the log-rank (Mantel-Cox) test. For Spontaneous seizures, multiple variables logistic regression modeling was computed to characterize odds ratio for an observed seizure event from the independent-variable main effects and two-way interactions (sex and genotype). Cohorts were not specifically powered to detect sex effects. Sample sizes were based on our previous studies using the same assays in mouse models of neurodevelopmental disorders. Significance levels were set at p<0.05.

### Data Availability

The data that support the findings of this study are available from the corresponding author, upon reasonable request. Codes used for MEA analysis can be found at the following link: https://github.com/BenShalomLab/MaxwellMEA-Toolkit-MATLAB/tree/V1

## Results

### Mouse model generation

The *KCNT1* gene sequences differ slightly between human and mouse: the altered amino acid in the G288S human *KCNT1* variant is at amino acid position 269 in the mouse protein (**Fig. 1**). Genome editing with CRISPR/Cas9 was used to make the mouse model: the WT mouse *Kcnt1* exon 11 codon (GGG) was replaced with AGC to create the missense mutation. This leads to glycine (G) being replaced by serine (S) at amino acid position 269 (G269S) in the mouse KCNT1 protein this new allele (*Kcnt1^em1Bryd^*) was nucleotide sequence verified. The mutation was generated on a C57BL/6J genetic background and was maintained by backcrossing heterozygous animals to B6/J and B6/NJ mice.

**Fig. 1.**
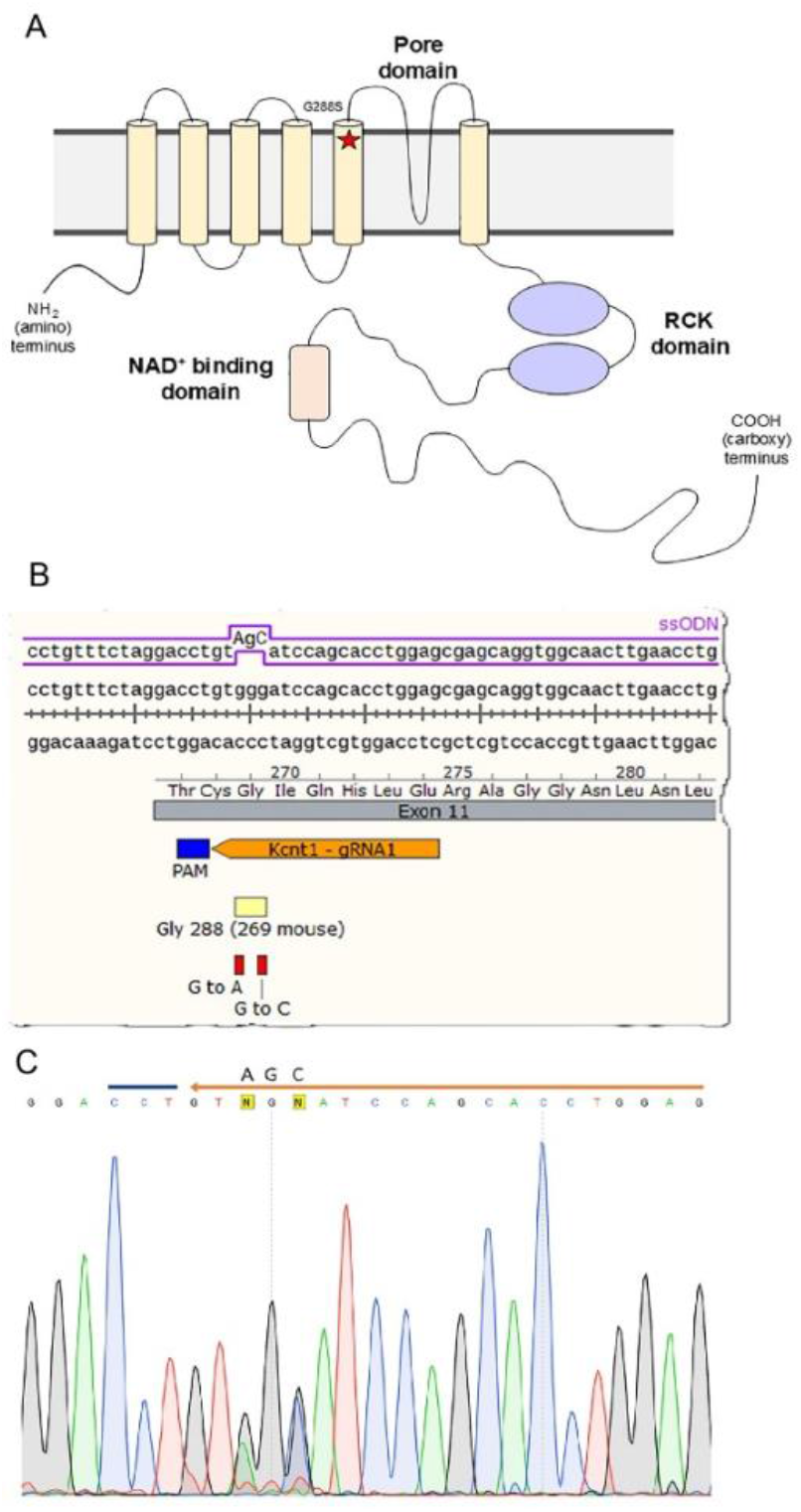
Generation of the *Kcnt1*^G269S^ mutant mouse model. (**A**) The KCNT1 protein consists of six hydrophobic transmembrane segments with a pore loop domain between the last two transmembrane segments. It also has a large intracellular carboxy-terminal region containing tandem RCK domains and an NAD+ binding domain. The mouse G269S (corresponding to human G288S) mutation, indicated by a red star, occurs within the channel pore. (**B**) The mouse model was engineered using CRISPR-Cas9 genome editing to replace the endogenous GGG (Gly) with AGC (Ser). (**C**) Confirmation of the engineered DNA base pair changes using Sanger sequencing. The heterozygous sample in the chromatogram shows overlapping G (black) and A (green) peaks for the G to A change and overlapping G (black) and C (blue) peaks for the G to C change.

### High-Density microelectrode array analysis reveals distinct network activity patterns in *Kcnt1*^+/G269S^ and *Kcnt1*^G269S/G269S^

To investigate the functional consequences of the *Kcnt1* G269S mutation at the network level, we employed high-density microelectrode array (HD-MEA) recordings of primary cortical neurons from wild-type (*Kcnt1^+/+^*), heterozygous (*Kcnt1^+/G269S^*), and homozygous (*Kcnt1^G269S/G269S^*) mice. The Maxwell Biosystems HD-MEA platform featuring 26,400 electrodes with 1,024 simultaneous recording channels enabled detailed spatial and temporal mapping of neuronal network activity patterns (**Fig. 2A**). Extracellular signals were recorded from multiple channels allowing for precise detection of action potentials across the neuronal population (**Fig. 2B, C**) from DIV 13-21, spanning the critical network formation and early maturation phases. To ensure comparable recording conditions across genotypes, we quantified the percentage of active electrodes: all three genotypes showed similar electrode activation rates (approximately 70-75%, **Fig. 2D**), indicating that the G269S mutation did not significantly affect neuronal viability or density during initial plating and attachment. However, despite similar electrode coverage, mean firing rates (DIV13-21) differed significantly between genotypes (**Fig. 2E**; F_(2,150)_=9.18, p=0.0002) in a gene-dosage dependent manner: *Kcnt1^G269S/G269S^* neurons had the highest mean firing rate (4.26±0.28 Hz), followed by *Kcnt1*^+/G269S^ (3.54 ± 0.15Hz) and *Kcnt1*^+/+^ neurons (2.81±0.16 Hz). Tukey’s post hoc tests revealed that *Kcnt1^G269S/G269S^* neurons fired significantly more than *Kcnt1*^+/G269S^ (p=0.0024) and *Kcnt1*^+/+^ neurons (p=0.0001); while *Kcnt1*^+/G269S^ neuronal firing rate was also elevated relative to *Kcnt1*^+/+^ (p=0.047).

**Fig. 2.**
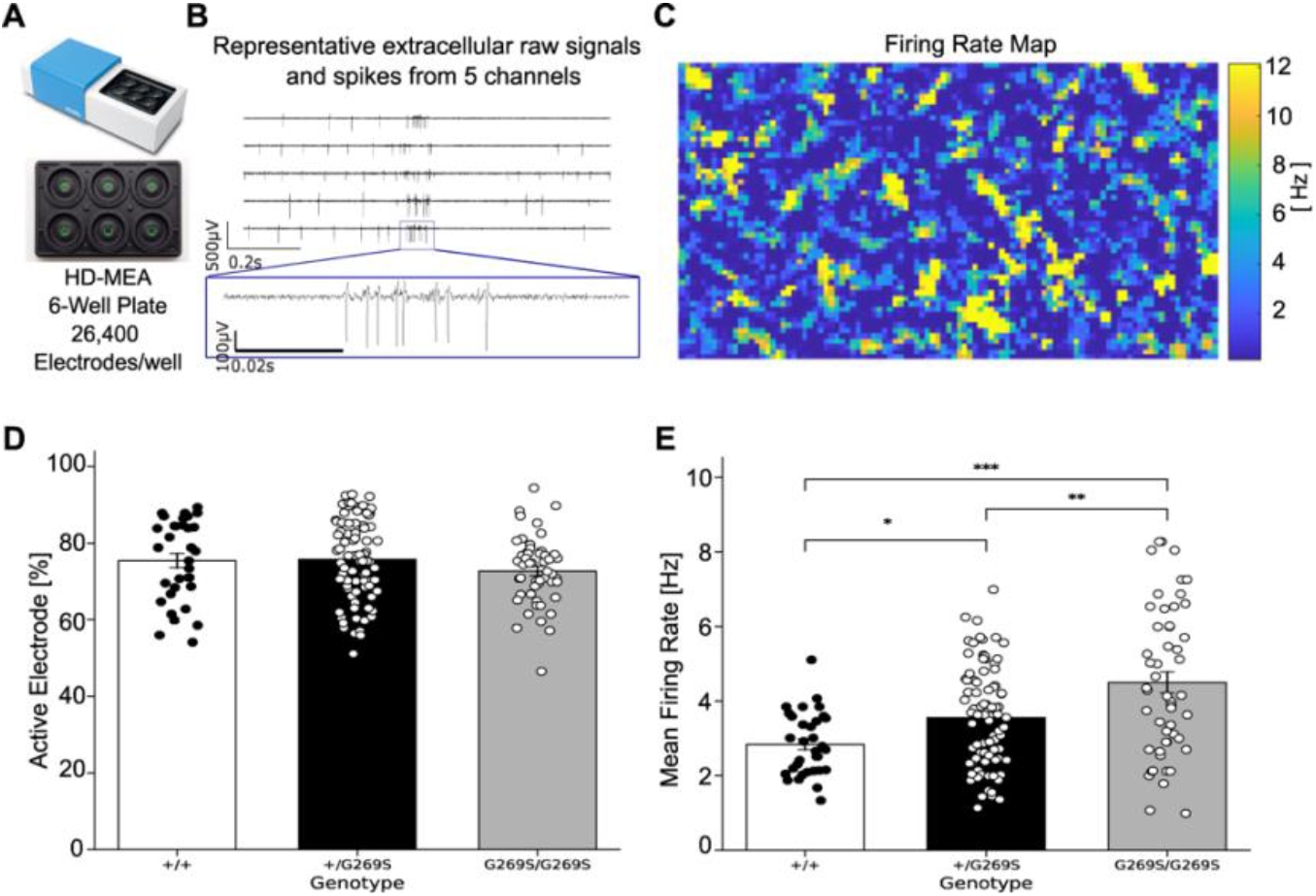
High-density MEA recordings reveal genotype-dependent differences in neuronal network activity. (**A**) Image of the high-density microelectrode array (HD-MEA) chip used in this study (26,400 electrodes; MaxTwo platform, Maxwell Biosystems). (**B**) Example of raw extracellular action potential (eAP) traces recorded from active electrodes. (**C**) Representative firing rate map from a cortical culture, showing spatial distribution of neuronal activity across the MEA surface. Warmer colors indicate higher firing rates (color scale: 2–12 Hz). (**D**) Percentage of active electrodes per well for wild-type *Kcnt1*^+/+^, heterozygous *Kcnt1*^+/G269S^, and homozygous *Kcnt1*^G269S/G269S,^ neuronal networks. (**E**) Quantification of mean firing rate (Hz) across genotypes. *Kcnt1*^G269S/G269S^ neurons fired significantly more than both *Kcnt1*^+/G269S^ and *Kcnt1^+/+^*. Bars represent mean ± SEM. Each point is one well (an independent culture) at one recording session; recordings from DIV13 to 21 were pooled (N = 153 well recordings from 55 independent wells; *Kcnt1*^+/+^ n = 10 wells [30 recordings], *Kcnt1*^+/G269S^ n = 30 wells [81 recordings], *Kcnt1*^G269S/G269S^ n = 15 wells [42 recordings]). In (**D**) no comparison reached significance. In (**E**), one-way ANOVA F(2,150) = 9.18, p =0.0002; Tukey post hoc: *Kcnt1*^G269S/G269S^ vs. *Kcnt1*^+/+^ p =0.0001, *Kcnt1*^G269S/G269S^ vs. *Kcnt1*^+/G269S^ p = 0.0024, *Kcnt1*^+/G269S^ vs. *Kcnt1*^+/+^ p = 0.047. p < 0.05 (*), p < 0.01 (**), p < 0.001 (***) by one-way ANOVA followed by Tukey post hoc test.

To confirm that our cultures contain both excitatory and inhibitory neurons and that these neurons are synaptically integrated in the network, we pharmacologically isolated GABAergic and glutamatergic signaling using HD-MEA recordings (**Fig. S1 and S2**). Gabazine, a GABA-A receptor antagonist, blocks inhibitory synaptic transmission. When we added gabazine (10µM) to HD-MEA chips and recorded for 600sec 1 hour later, network burst participation fraction, the fraction of active electrodes joining each detected network burst averaged across all bursts in the recording, increased significantly more in Treatment than in Vehicle Control wells in *Kcnt1*^+/G269S^ cultures (group × condition interaction, F(2,8)=25.58, p<0.001; Treatment change +0.16±0.06 vs. Vehicle change −0.02±0.01, unpaired t-test, p=0.007; **Fig. S2D**). Burst peak population firing rate, the population-wide firing rate at the single most active moment of each burst averaged across bursts, showed the same pattern (F(2,8)=28.94, p<0.001; Treatment change +49,800±6,273Hz vs. Vehicle change −5,732±4,108Hz, unpaired t-test, p<0.001; **Fig. S2F**), consistent with disinhibition of an active inhibitory network. Next, we applied AP5 (20 µM) and NBQX (20 µM) together. AP5 blocks NMDA receptors and NBQX blocks AMPA/kainate receptors; together they eliminate excitatory synaptic transmission. Blockade of glutamatergic signaling reduced the percentage of active electrodes, electrodes recording at least one spike during the 600 sec recording, to near zero in Treatment wells of both genotypes (*Kcnt1*^+/+^: Treatment change −87.9±7.2% vs. Vehicle change −2.3±2.5%, unpaired t-test, p<0.001; *Kcnt1*^+/G269S^: Treatment change −93.8±0.9% vs. Vehicle change 0.1±0.4%, unpaired t-test, p<0.001; **Fig. S2A, B**), with corresponding decreases in burst participation fraction (Treatment change −0.51±0.18 vs. Vehicle change 0.01±0.03, unpaired t-test, p=0.008) and burst peak population firing rate (Treatment change −102,215±41,060 Hz vs. Vehicle change −8,971±2,399Hz, unpaired t-test, p=0.017) in *Kcnt1*^+/G269S^ cultures. Together, these results confirm that our cultures contain synaptically connected excitatory and inhibitory neurons.

### Network development and bursting dynamics

Early network development corresponds to DIV 13 when synchronous activity begins to emerge and spread across the neuronal population,^36,37^ while DIV 27 represents the mature network phase characterized by established bursting patterns and stabilized connectivity.^38–40^ At DIV 13, we observed distinctive firing patterns across the three genotypes (**Fig. 3A-C** F_(2,52)_=6.515, p=0.003). During this network formation phase, *Kcnt1^G269S/G269S^* cultures displayed more frequent bursting activity compared to *Kcnt1*^+/+^ (Tukey’s post hoc p=0.002), with *Kcnt1^+/G269S^* networks showing intermediate burst frequency (Tukey*_Kcnt1+/+_*_vs*Kcnt1+/G269S*_ p=0.039; Tukey *_Kcnt1+/G269S_*_vs*Kcnt1G269S/G269S*_ p=0.199). By DIV 27, the differences in activity patterns became more pronounced but showed a different trajectory than at DIV 13 (**Fig. 3D-F** F_(2,41)_=13.97, p<0.0001). While *Kcnt1*^+/+^ and *Kcnt1^+/G269S^* cultures maintained consistent bursting patterns with increased synchronization characteristic of mature networks, *Kcnt1^G269S/G269S^* cultures exhibited markedly diminished burst amplitude and altered burst patterns (Tukey*_Kcnt1+/+_*_vs*Kcnt1G269S/G269S*_ p=0.0646; Tukey*_Kcnt1+/G269S_*_vs*Kcnt1G269S/G269S*_ p<0.0001). This unexpected reversal in the homozygous phenotype suggests potential network reorganization or adaptation following prolonged hyperexcitability during early development.

**Fig. 3.**
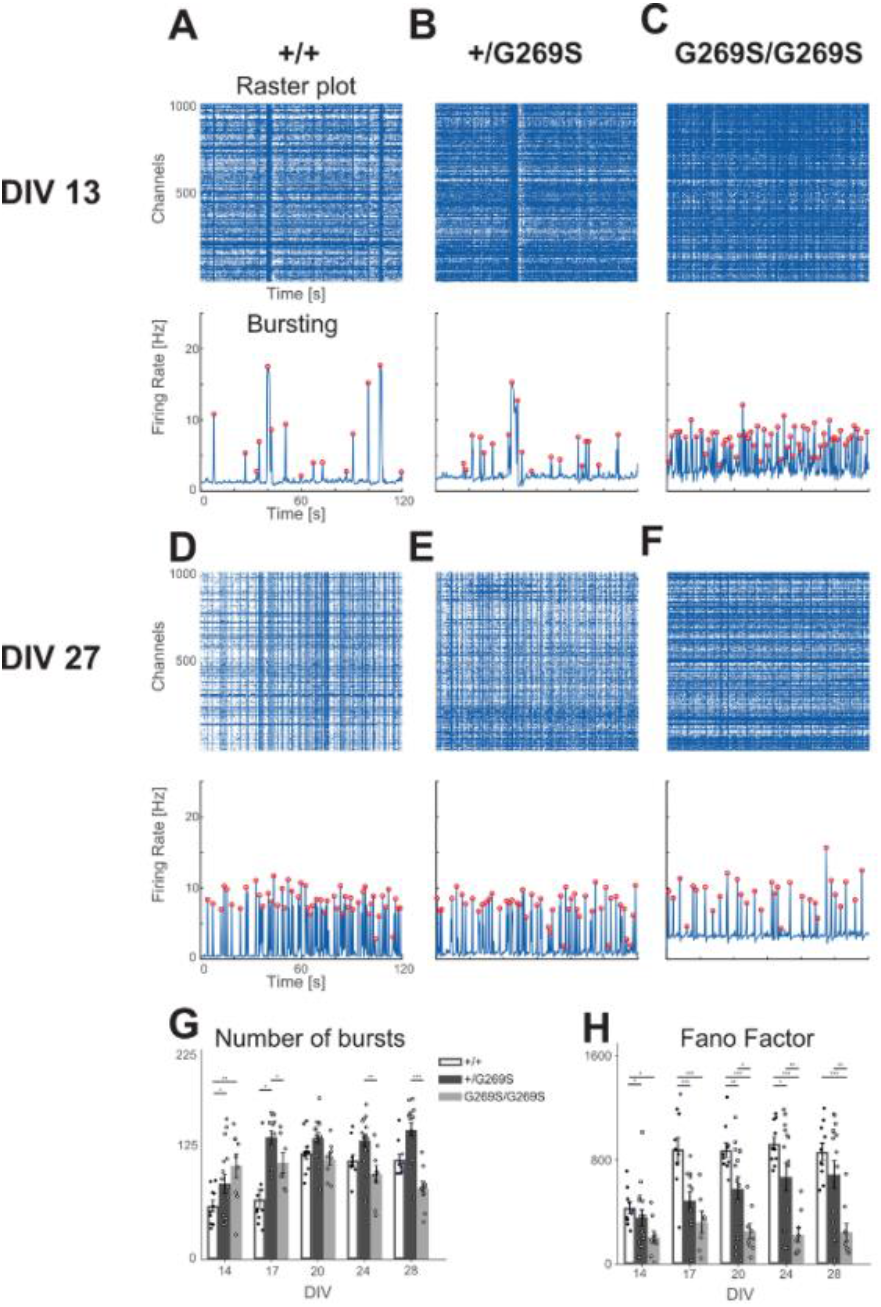
Developmental progression of burst dynamics in *Kcnt1*^+/+^, *Kcnt1*^+/G269S^, and *Kcnt1*^G269S/G269S^ in primary neuron culture. (**A–F**) Representative raster plots (top) and population firing rate traces (bottom) from wild-type (*Kcnt1^+/+^*, **A**, **D**), heterozygous (*Kcnt1^+/G269S^*, **B**, **E**), and homozygous (*Kcnt1^G269S/G269S^*, **C**, **F**) cortical networks recorded at DIV13 (**A–C**) and DIV27 (**D–F**). Each raster shows 120secs of activity across 1024 electrodes. Firing rate traces display synchronized burst events detected using a Gaussian kernel convolution; red dots indicate burst peaks. *Kcnt1^+/+^*and *Kcnt1^+/G269S^* networks display sparse, periodic bursts at DIV13 that become more frequent and structured by DIV27. In contrast, *Kcnt1^G269S/G269S^*networks show highly frequent and irregular bursting at DIV13, followed by diminished activity and synchrony at DIV27. (**G**) Quantification of burst count across developmental timepoints (DIV13–27). *Kcnt1^G269S/G269S^*networks display a biphasic pattern with early hyperexcitability (DIV13–17) and later hypoactivity, whereas *Kcnt1^+/G269S^* cultures maintain and increase burst frequency over the developmental time. (**H**) Fano factor, a measure of firing variability, is reduced in *Kcnt1*^G269S/G269S^ networks at all timepoints and in *Kcnt1*^+/G269S^ networks from DIV13 to DIV24 relative to *Kcnt1^+/+^*, indicating abnormally regularized activity. Bars represent mean ± SEM; each point is one well (an independent culture). One-way ANOVA at each DIV followed by Tukey post hoc tests (stars); a two-way ANOVA (genotype × DIV) tested effects across development. p < 0.05 (*), p < 0.01 (**), p < 0.001 (***). Recording days differed by one day between litters (see Methods), so adjacent DIVs were pooled; the x-axis labels 14, 20 and 28 denote DIV13/14, DIV20/21 and DIV27/28, reported as DIV13, 21 and 27 in the text and Supplemental Table 3. N per genotype and DIV, F values, and exact Tukey p values for every comparison are listed in **Supplemental Table 3**.

#### Temporal evolution of network hyperexcitability

Quantitative analysis of burst frequency across developmental timepoints (**Fig. 3G**; early window F(2,52)=9.55, p=0.0003; late window F(2,41)=10.99, p=0.0002) revealed a striking biphasic pattern in *Kcnt1^G269S/G269S^* cultures. Between DIV 13-17, *Kcnt1^G269S/G269S^* networks exhibited significantly higher burst counts (107.9±7.21) per recording, compared to *Kcnt1*^+/+^ (62.00±7.41, Tukey’s p=0.0002) reflecting initial hyperexcitability during network formation. However, bursting activity numerically declined in *Kcnt1^G269S/G269S^* networks (81.04±9.15) tending below *Kcnt1*^+/+^ levels (108.4±5.34) by DIV 24-27, though did not reach significance levels (Tukey’s p=0.078). In contrast, *Kcnt1^+/G269S^* cultures increased burst frequency over development, showing significantly elevated activity (94.68±4.74; Tukey’s p=0.0003) at DIV13-17 compared to *Kcnt1*^+/+^ and remaining numerically, but not significantly, elevated relative to *Kcnt1*^+/+^ levels (129.40±6.91 Tukey’s p=0.171) by DIV 24-27. Meanwhile, *Kcnt1^+/G269S^*cultures burst frequency was below *Kcnt1^G269S/G269S^* networks at DIV 13-17 (Tukey’s p=0.253), not significant, and rose above their levels by DIV 24-27 (Tukey’s p=0.0001). This indicates that the *Kcnt1^+/G269S^*confers a persistent and sustainable hyperexcitability phenotype that remains compatible with ongoing network function, unlike *Kcnt1^G269S/G269S^*.

Further analysis of the firing pattern variability was performed, using the Fano factor, the variance-to-mean ratio of spike counts, where a higher Fano factor reflects the healthy stochastic variability essential for proper circuit formation and refinement.^41^ *Kcnt1*^+/+^ networks maintained high Fano factor values (Fig. 3H, 656.31±56.30; early window F(2,52)=14.34, p<0.0001; late window F(2,41)=13.02, p<0.0001) In contrast, both *Kcnt1^G269S/G269S^* (303.11±38.52; Tukey’s p=0.004) and *Kcnt1^+/G269S^* networks (326.67±35.61; Tukey’s p<0.0001) showed significantly lower values during the early developmental window. While the Fano factor of *Kcnt1^+/G269S^* networks gradually trended higher toward *Kcnt1*^+/+^ levels by DIV 27 (Tukey’s p=0.122) but was not significant, *Kcnt1^G269S/G269S^* networks remained significantly reduced relative to *Kcnt1*^+/+^ (Tukey’s p<0.0001) and significantly diverged from *Kcnt1^+/G269S^* (Tukey’s p=0.0023), consistent with abnormally regularized firing pattern. This developmental trajectory (initial hyperexcitability followed by progressive reduction in network activity in *Kcnt1^G269S/G269S^* cultures, contrasted with sustained hyperexcitability in *Kcnt1^+/G269S^* cultures) suggests that the G269S GOF mutation affects network development in a gene-dosage dependent manner. Based on these electrophysiological findings, we decided to pursue only the *Kcnt1^+/G269S^* model for subsequent *in vivo* behavioral characterization.

#### Behavioral seizure identification and quantification, in the *Kcnt1^+/G269S^*mouse model by multiple methods

Spontaneous seizures were observed within an independent cohort during early life (**Table 1**). Within this cohort, a logistic regression model (AUC=0.65, p=0.17) identified a trend associated with genotype, sex, and an interaction (OR_GENOTYPE_=1.50, p=0.75; OR_SEX_=1.33, p=0.85; OR_INTERACTION_=2.18, p=0.66) indicating that *Kcnt1*^+/G269S^ males were associated with an elevated presentation of spontaneous seizure susceptibility, although this did not reach significance. Susceptibility to seizures early in life (post-natal day 21-23) was induced by hyperthermia, analogous to fever induced (febrile) seizures in Cohort 2 animals. Susceptibility was measured by probability to develop behavioral seizures, temperature to first seizure, and severity of seizure by a modified Febrile Racine score. *Kcnt1^+/G269S^* animals had a higher average Febrile Racine score than *Kcnt1*^+/+^ controls (**Fig. 4A**: t_(31)_=2.146, p=0.040) and a lower temperature threshold (**Fig. 4B**: t_(16)_=2.849, p=0.012). *Kcnt1^+/G269S^*animals also had an increased probability of developing a behavioral seizure (**Fig. 4C**: *χ*2=9.636, p=0.0019), over their *Kcnt1*^+/+^ counterparts. Importantly, a replicate cohort by a separate investigator reproduced the temperature threshold and seizure probability effects (**Fig. S3A-C**).

**Fig. 4:**
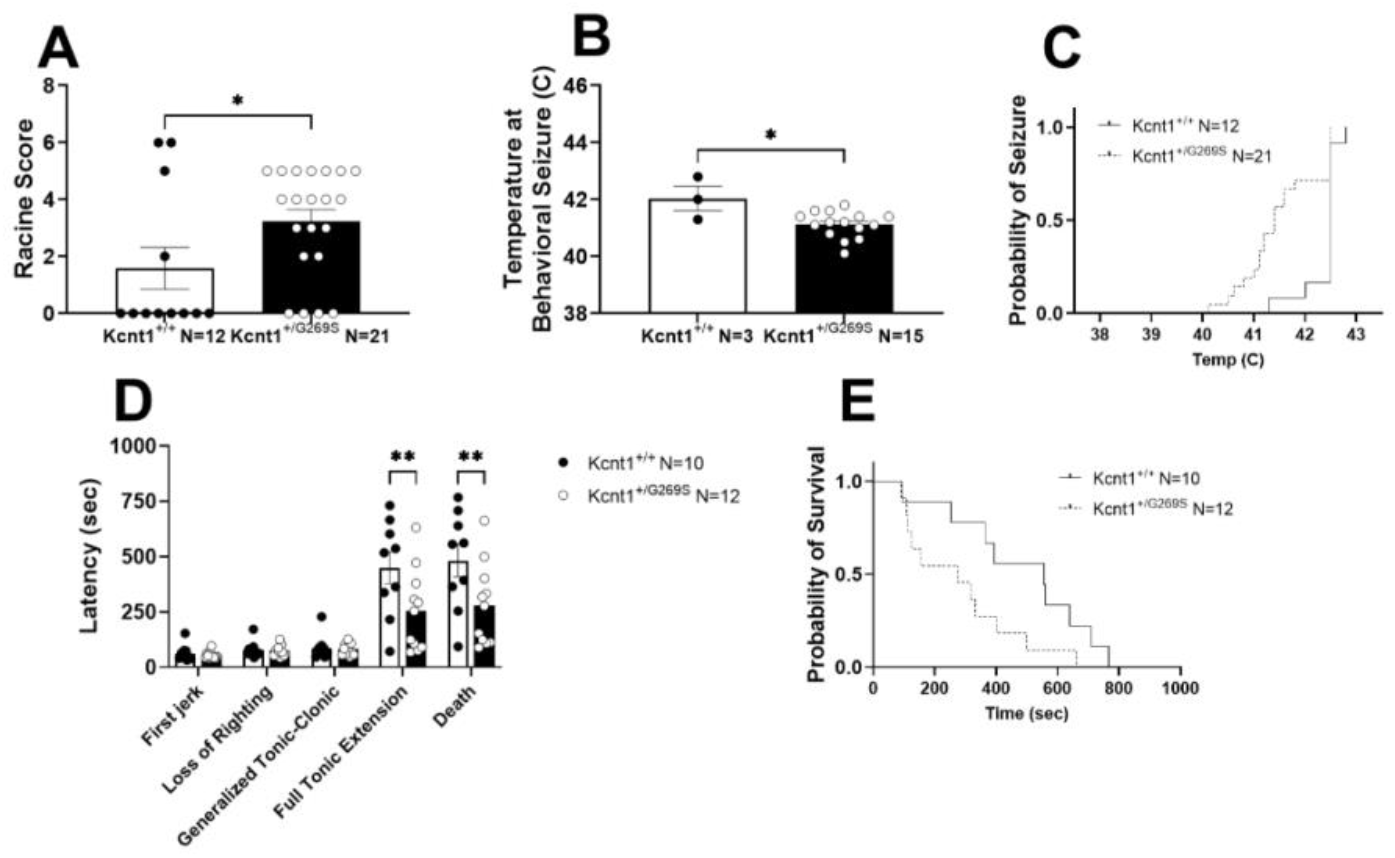
Seizure responsiveness to Hyperthermia-induction and the chemoconvulsant pentylenetetrazol in *Kcnt1*^+/G269S^ and *Kcnt1*^+/+^. (**A**) Seizure severity using modified Febrile Racine scale to Hyperthermia-induced seizure in N=33 mice; 5 female and 7 male *Kcnt1^+/+^;* 10 female and 11 male *Kcnt1^+/G269S^*(t_(31)_=2.146, p=0.040). (**B**) Average temperature to behavioral seizure (t_(16)_=2.849, p=0.012). (**C**) Probability of behavioral seizure as temperature is increased (χ2=9.636, p=0.0019). (**D**) Latency to each stage of a modified Racine scale in response to pentylenetetrazole (PTZ, i.p., 70 mg/kg) in N=22 mice; 5 female and 5 male *Kcnt1^+/+^;* 9 female and 3 male *Kcnt1^+/G269S^* (F_(4,76)TIME_=49.95, p<0.0001; F_(1,20)GENOTYPE_=4.96, p=0.038; F_(4,76)INTERACTION_=5.13, p=0.001; Bonferroni’s post hoc test: latency to full tonic extension p=0.004 and latency to death p=0.003). (**E**) Probability of survival following PTZ injection (χ2=3.878, p=0.049). Error bars indicate mean +/- SEM. * p<0.05 by either t-test (A,B) or repeated measures ANOVA followed by Bonferroni’s post hoc test (D) while probability of developing a behavioral seizure and probability of survival, were compared by chi-square test, and survival was analyzed by the log-rank (Mantel-Cox) test (C, E).

**Table 1.** Analysis of observed spontaneous seizures within an independent cohort during animal transit via multiple variables logistic regression model, which compared *Kcnt1^+/G269S^*female (N=18) and male (N=15) mice to *Kcnt1^+/+^* female (N=13) and male (N=10) control mice instances of sudden spontaneous epileptic activity. To characterize spontaneous seizure susceptibility, the odds probability ratio for each sex and genotypic variable was calculated where a seizure event was determined to be the dependent outcome variable. A potential, though non-significant (model AUC = 0.65, p = 0.17), trend where *Kcnt1^+/G269S^* mice (OR = 1.50, p=0.75) and male-sexed mice (OR = 1.33, p=0.85) were associated with an elevated presentation of spontaneous seizure susceptibility following transportation with reference to *Kcnt1^+/+^* and female mice, respectively. Notably, the interaction between genotype and sex (OR = 2.18, p=0.66) suggests a heightened vulnerability in *Kcnt1^+/G269S^* males. Animals were under temporary intrafacility room transfer from the vivarium at 5 and 6 weeks of age with handler precaution to maximally reduce cage disturbances.

| **Variable** | **Seizures / Total (n)** | **Seizure Rate (%)** | **Odds Ratio (OR)** | **95% Confidence Interval** | **Significance @ P<0.05** |
|---|---|---|---|---|---|
| Genotype [*Kcnt1<sup>+/G269S</sup>*] | 6/33 | 18.2% | 1.500 | 0.13 to 34.50 | 0.75 |
| Sex [Male] | 5/25 | 20.0% | 1.33 | 0.05 to 36.87 | 0.85 |
| *Kcnt1<sup>+/G269S</sup>* X Male Interaction | - | - | 2.18 | 0.05 to 93.67 | 0.66 |
| Model Performance | AUC: 0.65 | - | - | 0.44 to 0.8638 | 0.17 |

Adult behaviorally naive mice were measured for epileptic progression after induction with the chemoconvulsant pentylenetetrazol (PTZ, 70mg/kg, i.p.) *Kcnt1^+/G269S^* mice progressed through each seizure stage quicker than *Kcnt1^+/+^*mice (**Fig. 4D**: F_(4,76)TIME_=49.95, p<0.0001; F_(1,20)GENOTYPE_=4.96, p=0.038; F_(4,76)INTERACTION_=5.13, p=0.001). Bonferroni’s post hoc test revealed that *Kcnt1^+/G269S^* mice reached latency to full tonic extension (p=0.004), and latency to death (p=0.003) significantly faster than *Kcnt1^+/+^*mice, suggesting seizure susceptibility. Additionally, the probability of survival was decreased in *Kcnt1^+/G269S^* animals (**Fig. 4E**: *χ*2=3.878, p=0.049). Taken together, the *Kcnt1^+/G269S^* animals present spontaneous seizures, were highly sensitive to early life heat-induced seizures, as well as adult exposure to chemoconvulsive induced seizures.

### Exploratory locomotion and motor coordination in the *Kcnt1^+/G269S^*mouse model

*Kcnt1^+/G269S^* and *Kcnt1^+/+^* littermates did not differ in body weight, body length, head circumference or reflex development between PND 3 and 11 (**Supplemental Table 2**). Novel exploration in the open field was assessed between age, sex matched littermate controls, for effects resulting from *Kcnt1*^+/G269S^. Total activity across a 30min session illustrated an effect over time in both genotypes (**Fig. 5A**; F_TIME(5,148)_=7.706, p<0.0001), showing acclimation to the arena, as expected. Hyperactivity in the *Kcnt1*^+/G269S^ mice was observed when the 30min session was summed (**Fig. 5B**; t_(40)_=2.156, p=0.0372). Motor coordination and motor learning were measured using the accelerating rotarod. We observed significant reduction in latencies to fall in *Kcnt1*^+/G269S^ compared to *Kcnt1*^+/+^ age matched mice (**Fig. 5C**; F_GENOTYPE(1,42)_=6.79, p=0.0126) particularly on Day 1 and Day 2 by Tukey post hoc test (Tukey*_Kcnt1+/+_*_vs*Kcnt1+/G269S* DAY1_ p=0.002; Tukey*_Kcnt1+/+_*_vs*Kcnt1+/G269S* DAY2_ p=0.0151; Tukey*_Kcnt1+/+_*_vs*Kcnt1+/G269S* DAY3_ p=0.1517) indicating poor motor coordination. However, over the three training days we observed improved performance over time, (**Fig. 5C**; F_TIME(2,84)_=36.05, p<0.0001) in both genotypes illustrating intact motor learning.

**Fig. 5.**
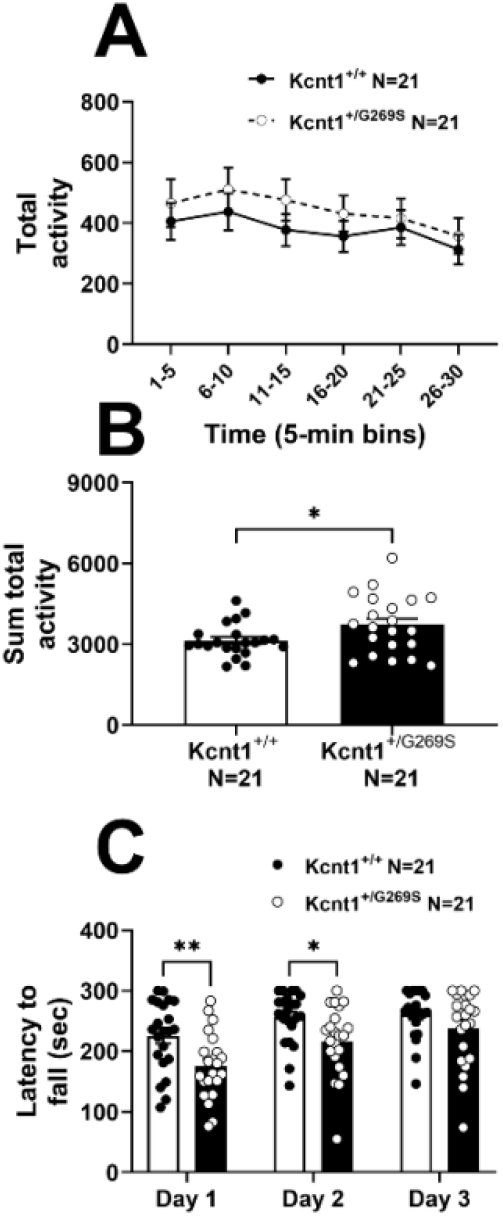
Elevated motor activity in a novel open field arena using the total activity parameter over a 30min period in *Kcnt1*^+/G269S^ vs *Kcnt1*^+/+^. Gross motor abilities were assessed using activity in a novel open field arena by: (**A**, **B**) total activity across a 30min session (t_(40)_=2.156, p=0.0372). In Panel A, data are shown in 5min bins (F_TIME(5,148)_=7.706, p<0.0001). (**C**) *Kcnt1*^+/G269S^ have poor motor coordination compared to *Kcnt1*^+/+^ controls using the standard accelerating rotarod paradigm on Days 1 and 2 of testing (F_GENOTYPE(1,42)_=6.79, p=0.0126 Tukey*_Kcnt1+/+_*_vs*Kcnt1+/G269S* DAY1_ p=0.002; Tukey*_Kcnt1+/+_*_vs*Kcnt1+/G269S* DAY2_ p=0.0151; Tukey*_Kcnt1+/+_*_vs*Kcnt1+/G269S* DAY3_ p=0.1517) with intact motor learning over time (F_TIME(2,84)_=36.05, p<0.0001). Error bars indicate mean +/- SEM. * p<0.05 by either a t-test (B) or repeated measures ANOVA followed by Tukey post hoc test (A, C). N=44 mice; 8 female and 14 male per genotype; 1 male from each genotype was removed from open field analysis following a Grubbs outlier test.

#### Cognitive ability assessed by two standardized assays in the *Kcnt1^+/G269S^*mouse model is intact

Cognitive ability was assessed in two tasks: novel object recognition (NOR) memory and fear conditioning. During the familiarization phase of the NOR task, Kcnt*1*^+/G269S^ mice explored the objects substantially less than the *Kcnt1*^+/+^ control mice (**Fig. S4A**: F_FAMILIARIZATION GENOTYPE(1,40)_=11.44, p=0.0016), with a mean investigation time in *Kcnt1*^+/+^ subjects of 51.79sec, while in *Kcnt1*^+/G269S^ the mean investigation time was 27.22sec. During the novel investigation phase, both genotypes spent greater time investigating the novel object, versus the familiar object (**Fig. S4B**: F_GENOTYPE(1,41)_=4.813, p=0.034 and Bonferroni’s post hoc test: *Kcnt1*^+/+^_FAMILIARvsNOVEL_ p=0.043; *Kcnt1*^+/G269S^_FAMILIARvsNOVEL_ p=0.0007). The discrimination index was calculated by dividing the difference in investigation time between the novel and familiar object by the total amount of exploration of both objects (**Fig. S4C**: t(40)=0.7566, p=0.45), clearly illustrating a lack of cognitive impairments in this assay. Cued and contextual fear conditioning illustrated a strong effect of training (learning from the tone-shock pairings (UCS-CS)) (**Fig. S4D**; F_TRAINING(1,40)_=107.8, p<0.0001). While a genotype by training interaction was observed, (**Fig. S4D**; F_TRAININGxGENOTYPE(1,40)_=4.951, p<0.04), there was no main effect of genotype (**Fig. S4D**; F_GENOTYPE(1,40)_=0.7672, p=0.3863), which is a key control metric, since no genotype effect across training allows for the investigation of differences in memory and recall. Contextual recall 24hrs after UCS-CS training did not differ between genotypes (**Fig. S4E**; t_(40)_=0.4610, p=0.6468). Cued recall, performed 48hrs after initial training, illustrated recall of the cued-shock pairing by high levels of freezing illustrated post-cue but not pre-cue in both *Kcnt1*^+/G269S^ and *Kcnt1*^+/+^ age matched mice (**Fig. S4F**; F_CUE(1,40)_=11.05, p=0.002).

#### Respiratory parameters identified erratic breathing and apnea in the *Kcnt1^+/G269S^*mouse model

Our laboratory was interested in defining respiratory metrics at baseline, given *KCNT1*-related epilepsy’s increased incidence of Sudden Unexpected Death in Epilepsy (SUDEP) and reports of apnea and oxygen desaturation during seizures in affected infants.^14,42,43^ Animals were assessed for 2hrs during the light cycle to quantify respiratory patterning and apnea occurrence in the whole-body plethysmography (WBP) chambers (**Fig. 6A)**. Analyses were stratified by sex because aged matched female mice are smaller than their male counterparts resulting in inherent sex differences in respiratory measurements. Percentage of time spent breathing normal was similar between *Kcnt1^+/G269S^*and *Kcnt1*^+/+^ females (**Fig. 6B**: t_(18)_=1.024 p=0.3194), whereas males exhibited decreased normal breathing (t_(20)_=3.237, p=0.0041). While, female *Kcnt1^+/G269S^*mice had increased erratic breathing (**Fig. 6C**: t_(18)_=2.635, p=0.0168) above their control counterparts, *Kcnt1^+/G269S^* males had a trend toward increased erratic breathing (**Fig. 6C**: t_(20)_=1.813, p=0.0842) that did not reach significance. The number of apnea events was elevated in *Kcnt1^+/G269S^*mice in males (**Fig. 6D**: t_(20)_=2.391, p=0.0258) and not in females (t_(18)_=0.4488, p=0.6589), compared to their respective sex matched *Kcnt1*^+/+^ controls. The duration of these apnea events were not different between female or male mice of either genotype (**Fig. 6E**: t_FEMALE(18)_=0.6885, p=0.4999; t_MALE(20)_=2.080, p=0.0506). The apnea events present in *Kcnt1^+/G269S^*male mice were categorized as Type 0 apneas, an extended period between an expiration and inspiration (**Fig. 6F**: t_FEMALE(18)_=0.4678, p=0.6455; t_MALE(20)_=2.830, p=0.0103). Taken together, this data may indicate that male *Kcnt1^+/G269S^* mice experience more respiratory difficulties than age matched females.

**Fig. 6:**
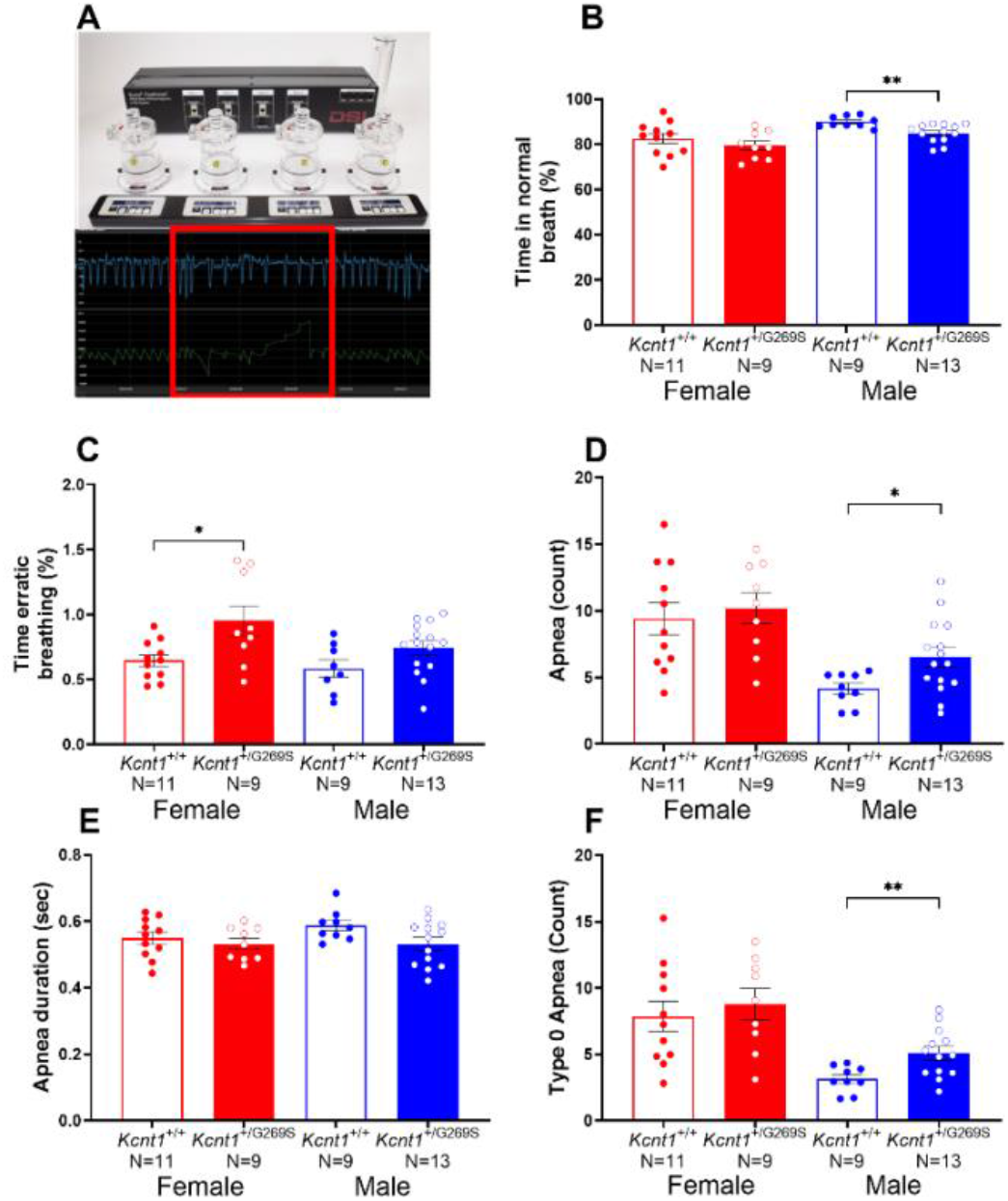
Respiratory apneas and breathing parameters assessed by whole body plethysmography (WBP) in *Kcnt1*^+/G269S^ and *Kcnt1* ^+/+^ mice. (**A**) Whole body plethysmography chambers, red box representative image of an apnea. (**B**) % of time breathing normal (t_FEMALE(18)_=1.024 p=0.3194; t_MALE(20)_=3.237, p=0.0041), (**C**) % of time breathing erratic (t_FEMALE(18)_=2.635, p=0.0168; t_MALE(20)_=1.813, p=0.0842), (**D**) number of apneas (t_FEMALE(18)_=0.4488, p=0.6589; t_MALE(20)_=2.391, p=0.0258), (**E**) apnea duration in seconds (t_FEMALE(18)_=0.6885, p=0.4999; t_MALE(20)_=2.080, p=0.0506)., (**F**) number of type 0 apneas (t_FEMALE(18)_=0.4678, p=0.6455; t_MALE(20)_=2.830, p=0.0103). Error bars indicate mean +/- SEM. * p<0.05 by t-test. N=42 mice; 11 female and 9 male *Kcnt1^+/+^;* 9 female and 13 male *Kcnt1^+/G269S^*.

## Discussion

Herein, we generated a novel experimental model for *KCNT1*-related epilepsy and channelopathies. This mouse model carries the mouse equivalent (G269S) of the human GOF variant G288S. CRISPR/Cas9 was used in C57BL/6J mice to replace the WT gene codon GGG with AGC, creating the missense mutation that substitutes glycine (G) to serine (S) at amino acid position 269 in the mouse KCNT1 protein. The new allele (*Kcnt1^em1Bryd^*) was nucleotide sequence verified, and the mice were maintained on C57BL6/NJ and were donated to the Mutant Mouse Resource and Research Center (MMRRC:075727-MU). Although the initial model was maintained on a C57BL/6NJ background, we backcrossed it onto C57BL/6J to align with existing models of developmental and epileptic encephalopathy (DEE) and rare genetic neurodevelopmental disorders (NDDs). However, we acknowledge the limitations of the C57BL/6J strain in seizure studies, as documented in prior literature.^20–22,44–46^ Given the reported differences in seizure susceptibility and responsiveness between sub strains, future experiments may return to the C57BL/6NJ background to better understand how genetic background influences phenotype, particularly of spontaneous recurrent seizures (SRS), seizure induction and potential of therapeutic testing^20,47^ Such comparisons are essential for evaluating the generalizability of findings and optimizing translational relevance.^48^

We performed functional characterization of primary cortical neurons from *Kcnt1*^+/+^, *Kcnt1*^+/G269S^, and *Kcnt1*^G269S/G269S^ mice *in vitro* revealing gene dose-dependent differences in excitability (**Fig. 2E**). Homozygous *Kcnt1*^G269S/G269S^ cultures showed an initial phase of hyperexcitability followed by a progressive decline in coordinated bursting, while heterozygous *Kcnt1*^+/G269S^ cultures maintained sustained hyperexcitability (**Fig. 3G**). Strikingly, both mutant genotypes exhibited a ∼50% reduction in firing pattern variability (**Fig. 3H**), indicating abnormally regularized activity. This loss of stochastic variability likely disrupts normal circuit development.^41^ Although *KCNT1* gain-of-function (GOF) variants, have been reported to affect excitatory and inhibitory neurons differently, with prior work describing preferential impairment of GABAergic interneurons^15,49^ our population-level HD-MEA recordings sample from a mixed network (**Fig. S1 and S2**) and cannot resolve the relative contribution of each cell type. Determining how the G269S variant differentially affects excitatory versus inhibitory neurons is an important direction for future work. The G269S mutation, located in the channel pore region, appears especially severe: homozygous cultures exhibit a progressive decline in coordinated bursting activity by DIV 28 (**Fig. 3D–G**), potentially reflecting homeostatic changes or network reorganization following sustained hyperexcitability. In contrast, the heterozygous model retains viable hyperexcitability, more accurately reflecting the human condition and offering a robust platform for therapeutic development and testing of currently available therapeutics.^50–52^

Next, we discovered that *Kcnt1^+/G269S^* mice not only exhibited spontaneous seizures in early life (**Table 1**), they were also susceptible to seizures following early life hyperthermia, a preclinical assay utilized as an analog for febrile seizures or seizures resulting from high temperature fevers, importantly humans with *KCNT1* variants experience early life seizures.^9^ Compared to age-matched control littermates, *Kcnt1^+/G269S^* mice seized at lower temperatures and were more likely to seize, and both effects replicated in a second cohort tested by a different investigator (**Fig. 4B, C**; **Fig. S3B, C**). Seizure severity on the modified febrile Racine scale was higher in the first cohort (**Fig. 4A**); the replicate showed the same direction without reaching significance (**Fig. S3A**). However, we do want to acknowledge the possible confound of using a brief anesthetic exposure prior to febrile seizure induction, this was used to prevent spontaneous seizures due to handling and insertion of the rectal probe. Additionally, behaviorally naïve adult *Kcnt1^+/G269S^*mice exposed to the chemoconvulsant pentylenetetrazol (PTZ, 70 mg/kg) were quicker to progress through the PTZ Racine scale compared to *Kcnt1^+/+^* controls (**Fig. 4D, E**). Additionally, we evaluated respiratory physiology using whole-body plethysmography (WBP), which was included given the clinical reports of asthma and pulmonary hemorrhage in *KCNT1*-related epilepsy,^53–56^ as well as the presence of apneas leading to disrupted sleep as a common occurrence in genetic NDDs, particularly when facial dysmorphic traits are evident.^57–59^ Male *Kcnt1^+/G269S^* mice exhibited increased apneas and females exhibited increased erratic breathing (**Fig. 6C, D, F**), which align with the clinical presentation. While *Kcnt1^+/G269S^* and *Kcnt1^+/+^* littermates were developmentally indistinguishable (**Supplemental Table 2**), *Kcnt1^+/G269S^*mice were hyperactive in the 30min open field arena (**Fig. 5A, B**) and lacked motor coordination on days 1 and 2 on the rotarod (**Fig. 5C**).

In summary, we report a novel *Kcnt1* G269S knock-in mouse model. Heterozygous *Kcnt1^+/G269S^* mice reproduce several features of *KCNT1*-related epilepsy, including spontaneous seizures in early life, heightened susceptibility to hyperthermia-induced and chemoconvulsant-induced seizures, hyperactivity, poor motor coordination and, in males, apneas, while developmental milestones, recognition memory and fear memory remain intact (**Supplemental Table 2**; **Fig. S4**). Because affected individuals carry a single pathogenic allele, the heterozygous model is the appropriate platform for *in vitro* and *in vivo* therapeutic testing; the homozygous *Kcnt1^G269S/G269S^* line lacks this construct validity.

Limited clinical success of KCNT1 channel blockers such as quinidine has been well documented.^14,50,51,60,61^ While cannabidiol (CBD), a compound derived from the cannabis plant, and ketogenic diet has had limited success in reducing seizure burden or severity in individuals with *KCNT1*-related epilepsy optimal treatment strategies are still under investigation.^62^ Additionally, reducing *KCNT1* gene expression using antisense oligonucleotides (ASO) has had some success in mouse models including in the homozygous P905L model^16^ and R455H^63^. Viral gene therapy and RNA interference approaches are also emerging: a divalent small interfering RNA produced durable seizure suppression in a preclinical mouse model of *KCNT1* epilepsy,^64^ and the current landscape of genetic and emerging therapies for *KCNT1*-related epilepsy has recently been reviewed.^65^

Future studies will incorporate long-term wireless EEG to assess cortical activity, including spiking, spike trains, sleep architecture, and spectral power density over multiple days. Cortical EEG is a critical translational biomarker and is routinely used by our lab and others,^26,35,66–68^ especially given the nuance of the C57BL6/J background strain.^69^ These analyses have the ability to illustrate the severity of this GOF variant and enable deeper in vivo phenotyping. These enhancements will contribute to a more comprehensive understanding of *KCNT1* variant effects and support improved patient stratification.

The G288S mutation has been identified in multiple unrelated patients with early-onset DEE.^14^ As previously noted, this variant does not correspond to a single epilepsy subtype,^9^ highlighting the importance of understanding how G288S/G269S variant modulates network excitability, thus insights from this novel in vivo model provide a tool for therapeutic development, and has potential to yield information relevant to the most affected individuals and inform stratification within the *KCNT1*-related epilepsy community. Groundwork in *KCNT1*-related epilepsy was outlined by many (for more see Supplemental Table 1). Highlights include the Wayne Frankel laboratory’s generation of the first ever *Kcnt1* mouse models, the knock-in (KI), human variants Y777H and Y796H. These experimental model systems illustrated motor cortex hyperexcitability, early-onset seizures, and hyperactivity in biallelic pathogenic KI/KI mice.^49,70^ In 2024, Wu et al.^15^ showed that the R455H variant increases both K_Na+_ and voltage-gated sodium currents in cortical neurons, enhancing the firing of excitatory neurons while suppressing that of inhibitory interneurons, together with upregulated NaV1.6 expression; K_Na+_ current increases were 2 to 4-fold in heterozygous and 4 to 6-fold in homozygous neurons. While previous findings in homozygous variant mice are significant, exhibiting the rigorous, robust effect sizes that are required for assessing in vivo therapeutic efficacy, unmet need existed for a construct valid single mono allelic KI variant in vivo model, since clinically, affected individuals with *KCNT1*-related epilepsy carry a single pathogenic allele. Herein, we have generated a tool to bridge the gap: our model presented herein has clear phenotypic excitability in primary cortical neurons, enhanced seizure sensitivity in early life as well as in adults, by hyperthermia and chemoconvulsant, respectively, as well as gross motor hyperactivity and poor motor coordination.

## Conclusions

We generated a heterozygous mouse model of a *KCNT1* pore-domain variant, *Kcnt1* G269S, which corresponds to the recurrent human G288S variant. A single mutant allele produced sustained hyperexcitability and hypersynchronous bursting in cortical networks, spontaneous seizures, heightened susceptibility to hyperthermia-induced and pentylenetetrazol-induced seizures, hyperactivity, poor motor coordination and, in males, apneas. These *in vitro* and *in vivo* measures provide quantitative endpoints for testing therapies, and the line is available to the community through the Mutant Mouse Resource and Research Center (MMRRC 075727-MU). Long-term EEG and sleep recordings will next define the neurophysiological basis of these phenotypes.

## Acknowledgements

The mouse model was generated by the University of Missouri Animal Modeling Core.

## Supplementary material

Supplementary material is available at *Brain Communications* online.

