## Supplement for "Neuronal Hyperexcitability and Seizure Susceptibility in a Novel *Kcnt1 G269S* Mouse Model"

### Supplemental Materials and Methods

#### Developmental Milestones

Developmental milestones were conducted by an experimenter blind to genotyped and were measured on postnatal days (PND) 3, 5, 9 and 11, similar to those previously described.[1] Physical characteristics of body weight and body length were assessed by a digital scale (grams) and the distance from the nose to edge of tail. Head width was measured using a digital caliper (cm). Reflexes tested were negative geotaxis, cliff avoidance, circle traverse and latency to righting, and fine motor abilities were measured by holding each pup and gently prodding the front paws with the wood end of a wooden sterile swab and measuring the ability of the pup to firmly grasp the stick. Lastly, the pups' ability to hang on a wire suspended over a softly padded cup was assessed by latency to fall, as our group has previously described for mice and rats.

#### Novel Object Recognition (NOR)

The NOR assay was conducted in a 30-lux room in an arena (P95 White, 41 cm, 41 cm w x 30 cm) The task consisted of 4 stages: (i) habituation to the opaque matte white arena for 30 min; (ii) a 10 min re-habituation session; (iii) a 10 min familiarization session during which mice were exposed to 2 identical objects, followed by a 60 min inter-trial-interval; and (iv) a 5 min recognition test, during which mice were exposed to an object from familiarization and one novel object in the same locations as in familiarization session, as previously described.[1–5] The objects were plastic toys: a small soft plastic orange safety cone and a hard plastic magnetic cone with ribbed sides and were counterbalanced across animals. Each session was recorded and visually tracked by contrast software Ethovision XT (Version 9.0, Noldus Information Technologies,

Leesburg, VA), and confirmed by hand scoring by an experimenter blind to genotypes. Object investigation was defined as time spent sniffing the object when the nose was oriented toward the object and the nose-object distance was 2cm or less. Recognition memory was defined as spending significantly more time interacting with the novel object compared to the familiar object by two-way repeated measures ANOVA followed by Bonferroni's post hoc test within each genotype. Side bias was confirmed by time spent interacting with the 2 identical objects during the familiarization session.

#### Contextual and cued fear conditioning

Associated learning was probed with delayed contextual and cued fear conditioning for wherein a novel context/cue predicts fear inducing stimulus, as previously published.[6,7] The fear conditioning chamber (Med Associates, St Albans, VT, USA) was enclosed in a sound-attenuating cubicle and interfaced with a PC installed with VideoFreeze software (version 1.12.0.0, Med Associates). The first environment was well lit (~100 lx) with a stainless-steel grid floor swabbed with almond odor cue (prepared from almond extract; McCormick; 1:100 dilution). Training on day 1 consisted of a 2 min acclimation period followed by three tone-shock pairings (80 dB tone, 30 sec duration; 0.5 mA foot shock, 1 sec duration; inter-shock interval 90 sec) and a 2.4 min period during which no stimuli were presented. Contextual fear conditioning was performed 24hr after training in the same context of grid floor, almond odor, and ~100 lx overhead lighting in the absence of the tone and foot shock. Cued fear conditioning was performed 48hr after training in a novel environment consisting of a white acrylic floor and black walls with a vanilla odor, overhead lighting was turned off. Animals were given a 3 min acclimation period followed by a 3 min presentation of the tone (80 dB). Cumulative time spent freezing in each condition was quantified by the VideoFreeze software.

### Supplemental Figures and Legends

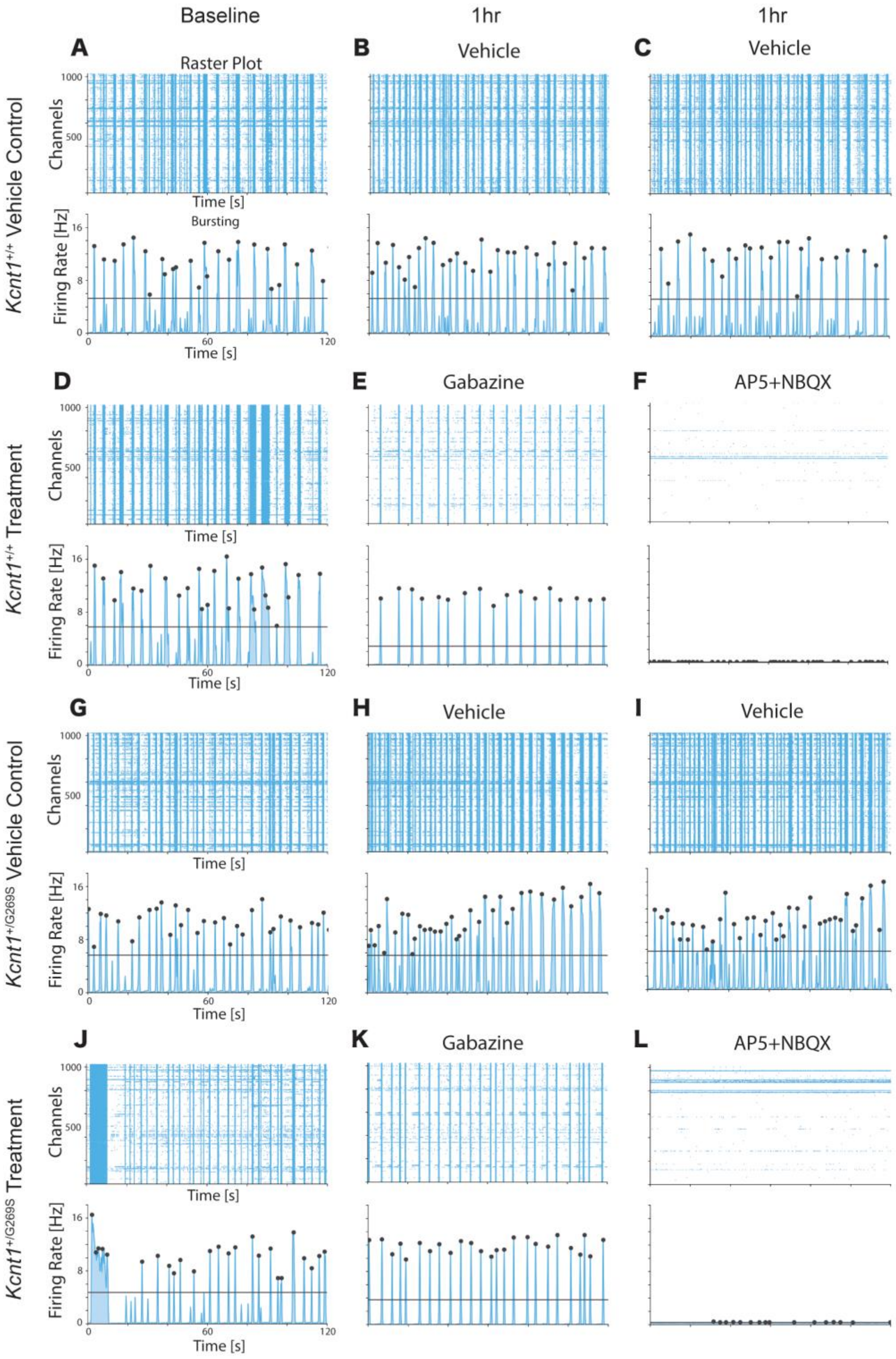


**Supplemental Figure 1: Pharmacological modulation of network burst activity in *Kcnt1*$^{+/+}$ and *Kcnt1*$^{+/G269S}$ cortical cultures, representative recordings.** (A–L) Representative raster plots (top) and population firing-rate traces (bottom) from a *Kcnt1*$^{+/+}$ Vehicle Control well (A–C), a *Kcnt1*$^{+/+}$ Treatment well (D–F), a *Kcnt1*$^{+/G269S}$ Vehicle Control well (G–I), and a *Kcnt1*$^{+/G269S}$ Treatment well (J–L), recorded at Baseline, 1 hr after the first addition, and 1 hr after the second addition (2 hr after Baseline). Each raster shows 120 s of activity across 1024 channels. Vehicle Control wells received water at both timepoints (B, C, H, I); Treatment wells received gabazine (10 μM) at the first timepoint (E, K) and AP5 (20 μM) plus NBQX (20 μM) at the second (F, L). Firing-rate traces display population activity with burst peaks marked by black dots and the burst-detection threshold indicated by a horizontal line. Vehicle Control wells in both genotypes maintain consistent bursting across all three recordings; Treatment wells show increased burst synchrony following gabazine (E, K) and a near-complete loss of network activity following AP5+NBQX (F, L). Quantification of these recordings is shown in Supplemental Figure 2.

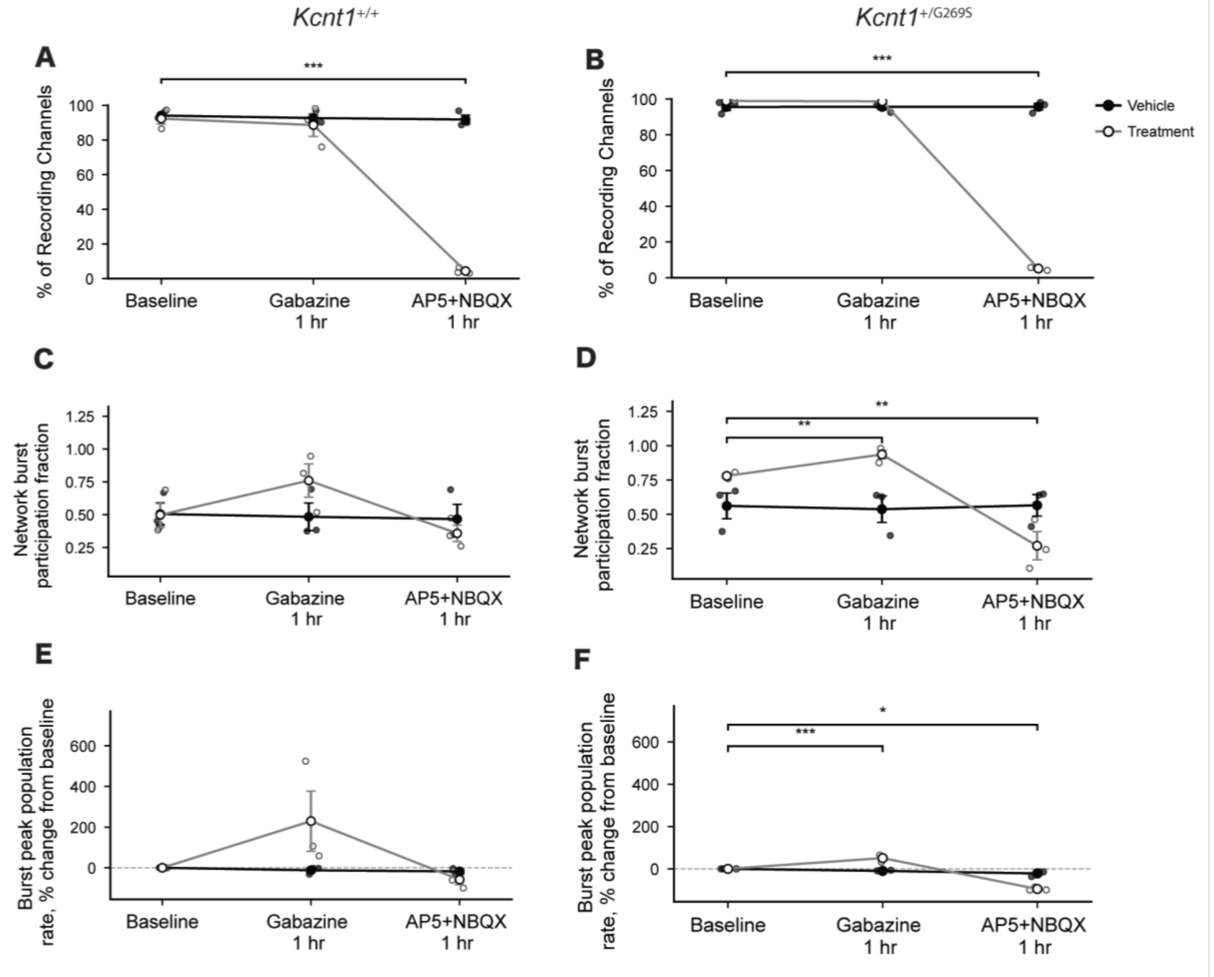


**Supplemental Figure 2: Pharmacological modulation of network burst activity in *Kcnt1*$^{+/+}$ and *Kcnt1*$^{+/G269S}$ cortical cultures, quantification.** Summary metrics for the wells shown in Supplemental Figure 1. (A, B) Percentage of recording channels with at least one spike during the recording in *Kcnt1*$^{+/+}$ (A) and *Kcnt1*$^{+/G269S}$ (B) wells. Vehicle Control (black) wells remain stable across all three conditions in both genotypes; Treatment (grey) wells are indistinguishable from Vehicle Control at Baseline and after gabazine but collapse to near-zero activity following AP5+NBQX in both genotypes (***, Baseline vs. AP5+NBQX 1 hr). (C, D) Network burst participation fraction, the mean fraction of active electrodes joining each detected network burst, in *Kcnt1*$^{+/+}$ (C) and *Kcnt1*$^{+/G269S}$ (D) wells. In *Kcnt1*$^{+/G269S}$, Treatment wells show a significant increase in participation fraction following gabazine and a significant decrease following AP5+NBQX relative to Baseline (**, both comparisons); the same trend in *Kcnt1*$^{+/+}$ does not reach statistical significance. (E, F) Burst peak population firing rate, expressed as percent change from each well's own Baseline, in *Kcnt1*$^{+/+}$ (E) and *Kcnt1*$^{+/G269S}$ (F) wells. In *Kcnt1*$^{+/G269S}$, Treatment wells show a significant increase following gabazine (***) and a significant decrease following AP5+NBQX (*) relative to Baseline; the same trend in *Kcnt1*$^{+/+}$ does not reach statistical significance. For A–F, n = 3 wells per group; points represent individual wells and error bars indicate SEM. Statistics: mixed-design ANOVA (group × condition; group as a between-subjects factor and condition as a within-subjects factor) followed by post hoc unpaired t-tests comparing each well's change from Baseline between Treatment and Vehicle Control groups; $p < 0.05$ (*), $p < 0.01$ (**), $p < 0.001$ (***); unmarked comparisons did not reach significance; p values were not corrected for multiple comparisons. Because the burst-detection threshold scales with the number of active electrodes (0.03 × active electrodes), it falls to about one electrode after AP5+NBQX, so burst-derived metrics (C–F) at the AP5+NBQX timepoint should be interpreted with caution; the effect of glutamatergic blockade is best read from panels A and B, which do not depend on burst detection.

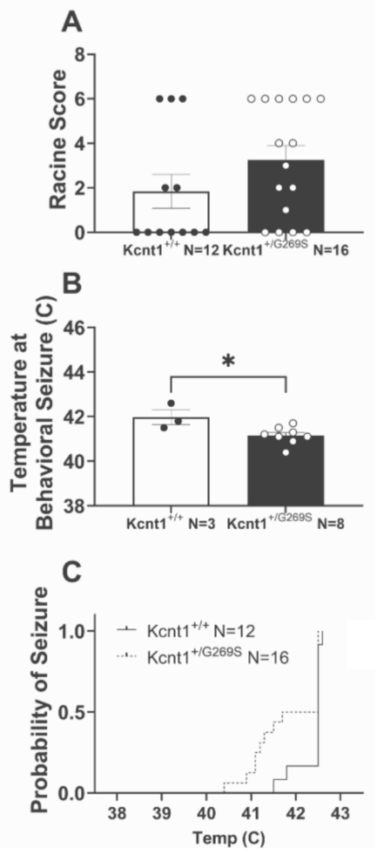


**Supplemental Figure 3: Seizure responsiveness to hyperthermia-induction in replicate Cohort** (A.) Seizure severity using modified Febrile Racine scale to Hyperthermia-induced seizure did not differ significantly between *Kcnt1*$^{+/G269S}$ (N=16) and *Kcnt1*$^{+/+}$ (N=12) controls (unpaired t-test, $t_{(26)}=1.44$, p=0.161) (B) Average temperature to behavioral seizure was decreased in *Kcnt1*$^{+/G269S}$ mice (unpaired t-test, $t_{(9)}=2.75$, p=0.022). (C) Probability of behavioral seizure as temperature is increased was increased in *Kcnt1*$^{+/G269S}$ mice ($\chi 2=4.52$, p=0.034). Error bars indicate mean +/- SEM. * p<0.05 by unpaired t-test (B) or chi-square test (C). Each point is one mouse.

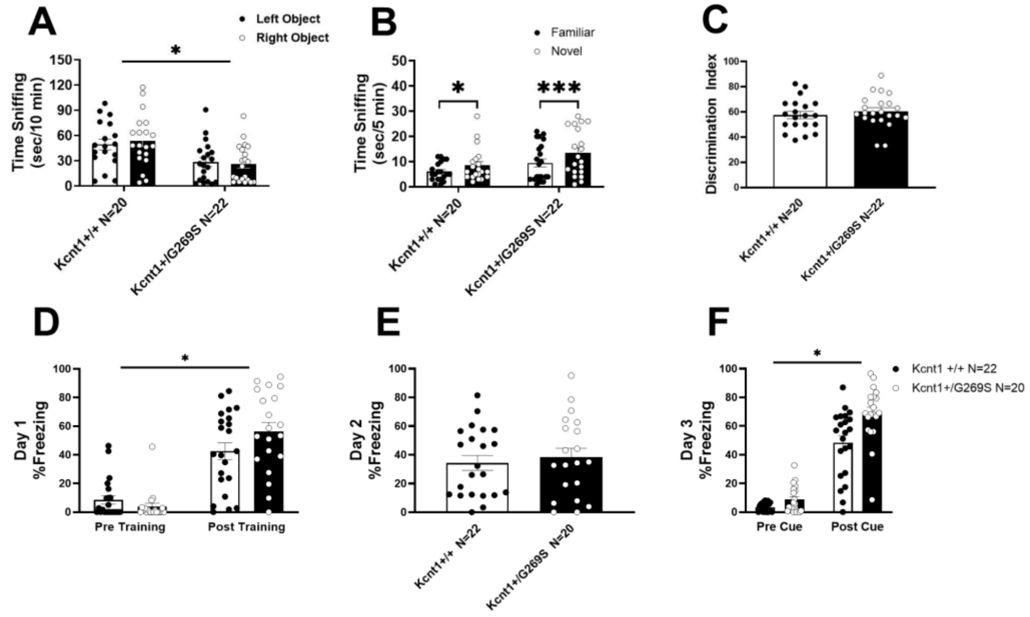


**Supplemental Figure 4: Learning and memory was intact in *Kcnt1*$^{+/G269S}$ mice.** Learning and memory was probed with the novel object recognition task wherein animals (*Kcnt1*$^{+/+}$ (N=20 after exclusion of 1 animal with 0 exploration and 1 animal with 95% exploration of one object) *Kcnt1*$^{+/G269S}$ (N=22) **)** explore 2 identical objects during the familiarization phase (A; $F_{FAMILIARIZATION\ GENOTYPE(1,40)}$=11.44, p=0.0016) followed by a 1 hour delay before testing for recognition memory (B; $F_{GENOTYPE(1,41)}$=4.813, p=0.034 and Bonferroni's post hoc test: *Kcnt1*$^{+/+}_{FAMILIARvsNOVEL}$ p=0.043; *Kcnt1*$^{+/G269S}_{FAMILIARvsNOVEL}$ p=0.0007). With no differences in the discrimination index between *Kcnt1*$^{+/G269S}$ and *Kcnt1*$^{+/+}$ controls (C; $t_{(40)}$=0.7566, p=0.45). During fear conditioning training there were no differences in response to the CS in either genotype on Day 1 (D *Kcnt1*$^{+/+}$ (N=22) and *Kcnt1*$^{+/G269S}$ (N=20) $F_{TRAINING(1,40)}$=107.8, p<0.0001; $F_{GENOTYPE(1,40)}$=0.7672, p=0.3863; $F_{TRAININGxGENOTYPE(1,40)}$=4.951, p<0.04). Similar levels of freezing were seen to the context probe on Day 2 (E) in both genotypes ($t_{(40)}$=0.4610, p=0.6468). On Day 3 freezing in a novel context to the cue was increased in both genotypes (F; $F_{CUE(1,40)}$=11.05, p=0.002). Error bars indicate mean +/- SEM. * p<0.05 by two-way repeated measures ANOVAs followed by Bonferroni's post hoc tests for selected group comparisons or unpaired t-tests as appropriate. Each point is one mouse.

#### Supplemental Table 1:

| Reference | Model | Genotypes Tested | Background Strain | Phenotypes found |
|---|---|---|---|---|
| Quraishi et al 2020 | Mice | Generated R474H variant mutation in mice *Kcnt1*$^{+/R455H}$<br><br>Knock-out mice (*Kcnt1*$^{-/-}$)<br><br>WTs (Kcnt1$^{+/+}$) were from a parallel colony of Sv/129 backcrossed with C57BL/6NJ | C57BL/6J and backcrossed for 4-5 generations<br><br>Sv/129 and backcrossed with C57BL/6NJ for 2 generations | • Knock out mice were hypoactive in open field and on the rotarod<br>• No seizure susceptibility<br>• *Kcnt1*$^{R455H/R455H}$ were embryonic lethal<br>• *Kcnt1*$^{+/R455H}$ mice<br> ○ Increased spontaneous seizures in EEG<br> ○ Decreased threshold for PTZ (60mg/kg)<br> ○ No motor differences in the open field or rotarod |
| Shore et al 2020 | Mice | Y796H (mouse Y777H) | C57BL/6NJ | • GABAergic neurons hyperexcitability and hyper-synchronicity<br>• Epileptiform activity was found in HOM mice via Video-EEG |
| Burbano et al 2022 | Mice | P924L (mouse P905L) | C57BL/6J | • HET mice had no seizure phenotypes<br>• HOM mice had spontaneous tonic-clonic seizures as early as P18, shortened life span (median survival 43 days), hyperactivity in the open field, and anxiety-like behavior.<br>• ASO treatment reduced seizures and prolonged lifespan of HOM |
| Gertler et al 2022 | Mice | L457F | C57BL/6J | • No significant differences between genotypes on Rotarod, Zero maze, Open Field, home cage<br>• No differences by genotype to seizure susceptibility by flurothyl (20ul/min) |

| | | | | |
|---|---|---|---|---|
| | | | | • KI/+ and KI/KI mice were susceptible to both Kainate (30mg/kg) and Pentylenetrazole (60mg/kg) induction |
| Tobin and Westin 2023 | Mice | Y796H (mouse Y777H) crossed with Snap25-GCaMP6s | C57BL/6NJ | • Widefield Ca2+ imaging in HOM and WT mice showed no differences in mesoscale interictal activity |
| Wu et al 2024 | Mouse primary neurons and slice | R474H (mouse R455H) | C57BL/6J | • Increased firing of excitatory neurons<br>• Suppressed firing of inhibitory neurons |
| Shore et al 2024 | Mouse primary neurons | Y796H (mouse Y777H) | C57BL/6NJ | • No effects on glutamatergic or VIP neuron function.<br>• SST neurons became hypoexcitable whereas PV neurons became hyperexcitable |
| Sun et al 2024 | Mice | Knock-out mice ($Kcnt1^{-/-}$) | C57BL/6J and backcrossed to C57BL/6NJ for >6 generations | • Complete loss of hippocampal LTP and LTD<br>• Reduced NMDA receptor signaling<br>• Localized protein changes |
| | | R455H/+ | C57BL/6J | • Some mild protein changes reversed with ASO treatment |
| Gou et al 2025 | Mice | Y796H (mouse Y777H) | C57BL/6 | • Potassium channel blockers, tipepidine and hydroquinine, were effective at blocking spontaneous seizures in Y777H HOM mice |
| Wang et al 2025 | Mice | Y796H (mouse Y777H) | on background strain C57BL/6NJ, but backcrossed to C57BL/6J | • Spontaneous seizures in HOM mice occur during NREM sleep and suppress REM sleep. |

**Supplemental Table 1: Previously generated mouse models of disrupted *Kcnt1* expression.** Summary of genotypes, background strains, and phenotypes reported for existing *Kcnt1* mouse lines (Quraishi et al. 2020; Shore et al. 2020; Burbano et al. 2022; Gertler et al. 2022; Tobin and Westin 2023; Wu et al. 2024; Shore et al. 2024; Sun et al. 2024; Gou et al. 2025; Wang et al. 2025).

| Milestones | Domain | Genotype | PND 3 | PND 5 | PND 9 | PND 11 | F Value | Significance @ P<0.05 |
|---|---|---|---|---|---|---|---|---|
| | | | Mean +/- SE | Mean +/- SE | Mean +/- SE | Mean +/- SE | | |
| Weight (g) | Physical | *Kcnt1+/+* | 1.64+/-0.07 | 2.40+/-0.09 | 3.90+/-0.12 | 4.70+/-0.15 | F (1, 23) = 0.79 | no, P=0.38 |
| | | *Kcnt1+/G269S* | 1.74+/-0.09 | 2.49+/-0.12 | 4.17+/-0.135 | 4.92+/-0.14 | | |
| Total length (mm) | Physical | *Kcnt1+/+* | 45+/-0.52 | 52.40+/-0.66 | 66.4+/-1.11 | 72.40+/-1.25 | F (1, 23) = 0.04 | no, P=0.85 |
| | | *Kcnt1+/G269S* | 45.71+/-1.07 | 51.86+/-1.43 | 65.86+/-1.09 | 71.93+/-1.07 | | |
| Head circumference (mm) | Physical | *Kcnt1+/+* | 8.17+/-0.18 | 9.00+/-0.19 | 10.30+/-0.19 | 11.40+/-0.20 | F (1, 23) = 0.30 | no, P=0.59 |
| | | *Kcnt1+/G269S* | 8.07+/-0.13 | 9.07+/-0.20 | 10.64+/-0.20 | 11.43+/-0.14 | | |
| Circle transverse (s) | Reflex | *Kcnt1+/+* | 30+/-0 | 29.60+/-0.45 | 26.3+/-2.06 | 21.60+/-2.88 | F (1, 23) = 0.07 | no, P=0.79 |
| | | *Kcnt1+/G269S* | 30+/-0 | 30+/-0 | 28.90+/-1.00 | 19.91+/-2.92 | | |
| Negative Geotaxis (s) | Reflex | *Kcnt1+/+* | 28.40+/-1.39 | 21.9+/-3.11 | 11.5+/-1.69 | 13.30+/-2.08 | F (1, 23) = 1.51 | no, P=0.23 |
| | | *Kcnt1+/G269S* | 25.48+/-2.15 | 21.40+/-2.60 | 11.7+/-2.15 | 8.23+/-1.35 | | |
| Righting (s) | Reflex | *Kcnt1+/+* | 27.15+/-2.13 | 27.7+/-2.32 | 6.70+/-3.33 | 2.90+/-1.40 | F (1, 23) = 2.17 | no, P=0.15 |
| | | *Kcnt1+/G269S* | 25.94+/-2.24 | 24.47+/-2.25 | 1.59+/-0.21 | 2.30+/-1.28 | | |
| Cliff aversion (s) | Reflex | *Kcnt1+/+* | 30+/-0 | 17.40+/-3.90 | 1.30+/-0.10 | 1.00+/-0.10 | F (1, 23) = 0.01 | no, P=0.90 |
| | | *Kcnt1+/G269S* | 26.22+/-2.6 | 18.36+/-3.70 | 3.52+/-2.00 | 0.89+/-0.10 | | |
| Hindlimb hang (s) | Reflex | *Kcnt1+/+* | 12.80+/-1.59 | 17.0+/-2.34 | 9.30+/-2.27 | 12.00+/-2.57 | F (1, 23) = 3.85 | no, P=0.06 |
| | | *Kcnt1+/G269S* | 14.10+/-1.92 | 10.14+/-1.87 | 8.41+/-0.96 | 9.12+/-1.61 | | |

**Supplemental Table 2: Developmental milestones.** Physical metrics and neurological reflexes on PND 3, 5, 9 and 11 compared between *Kcnt1*$^{+/G269S}$ (N = 14) and *Kcnt1*$^{+/+}$ (N = 11) sex- and age-matched littermates by repeated measures ANOVA; no developmental delays were detected.

| DIV | ANOVA F(df) | ANOVA p | Kcnt1+/+ Mean±SEM (n) | Kcnt1+/G269S Mean±SEM (n) | Kcnt1G269S/G269S Mean±SEM (n) | *Kcnt1+/G269S* vs *Kcnt1G269S/G269S* p-adj | *Kcnt1+/G269S* vs *Kcnt1+/+* p-adj | *Kcnt1G269S* vs *Kcnt1+/+* p-adj |
|---|---|---|---|---|---|---|---|---|
| **13** | F(2,52)=6.51 | **0.0030** ** | 58.5±6.2 (10) | 85.3±5.5 (30) | 101.3±8.3 (15) | 0.1991 | **0.0385** * | **0.0020** ** |
| **17** | F(2,52)=4.83 | **0.0119** * | 65.5±10.5 (10) | 104.1±7.7 (30) | 114.5±10.3 (15) | 0.6894 | **0.0295** * | **0.0115** * |
| **21** | F(2,41)=2.17 | 0.1276 | 116.1±6.8 (10) | 120.8±6.2 (20) | 100.6±8.7 (14) | 0.1133 | 0.9055 | 0.3883 |
| **24** | F(2,41)=6.43 | **0.0037** ** | 107.9±6.4 (10) | 122.7±7.2 (20) | 83.1±9.9 (14) | **0.0025** ** | 0.4573 | 0.1544 |
| **27** | F(2,41)=13.97 | **<0.0001** *** | 108.8±6.0 (10) | 136.1±7.3 (20) | 79.0±9.3 (14) | **<0.0001** *** | 0.0717 | 0.0646 |

**Supplemental Table 3: Statistics for Burst Count in Fig. 3G.**

Number of wells per genotype at each DIV, one-way ANOVA F values at each DIV, two-way ANOVA (genotype × DIV) results, and exact Tukey post hoc p values for every pairwise comparison shown in Fig. 3G (burst count). Two-way ANOVA (genotype × DIV): Genotype $F_{(2,227)}=9.77$, p=8.5e-5; DIV $F_{(4,227)}=7.27$, p=1.6e-5; Genotype × DIV $F_{(8,227)}=5.79$, p=1.0e-6. One-way ANOVA at each DIV followed by Tukey post hoc tests; a two-way ANOVA (genotype × DIV) tested effects across days in vitro. Recording days differed by one day between litters, so adjacent DIVs were pooled; DIV13, 21 and 27 denote DIV13/14, DIV20/21 and DIV27/28 (x-axis labels 14, 20 and 28 in Fig. 3). N = independent wells (cultures). Shaded cells: p<0.05. * p<0.05, ** p<0.01, *** p<0.001.

| DIV | ANOVA F(df) | ANOVA p | Kcnt1+/+ Mean±SEM (n) | Kcnt1+/G269S Mean±SEM (n) | Kcnt1G269S/G269S Mean±SEM (n) | *Kcnt1+/G269S* vs *Kcnt1G269S* p-adj | *Kcnt1+/G269S* vs *Kcnt1+/+* p-adj | *Kcnt1G269S* vs *Kcnt1+/+* p-adj |
|---|---|---|---|---|---|---|---|---|
| **13** | F(2,52)=4.15 | **0.0213 *** | 432.8±44.0 (10) | 269.0±37.9 (30) | 232.2±34.6 (15) | 0.7937 | **0.0408 *** | **0.0226 *** |
| **17** | F(2,52)=20.16 | **<0.0001 ***** | 879.9±93.8 (10) | 384.3±39.3 (30) | 374.0±48.5 (15) | 0.9885 | **<0.0001 ***** | **<0.0001 ***** |
| **21** | F(2,41)=13.44 | **<0.0001 ***** | 875.8±56.9 (10) | 545.4±73.5 (20) | 297.2±59.0 (14) | **0.0306 *** | **0.0081 **** | **<0.0001 ***** |
| **24** | F(2,41)=15.36 | **<0.0001 ***** | 925.4±51.5 (10) | 638.9±82.0 (20) | 258.9±67.1 (14) | **0.0019 **** | **0.0434 *** | **<0.0001 ***** |
| **27** | F(2,41)=10.59 | **0.0002 ***** | 862.5±67.7 (10) | 668.5±86.5 (20) | 272.7±79.5 (14) | **0.0035 **** | 0.2884 | **0.0003 ***** |

**Supplemental Table 4: Statistics for Fano Factor in Fig. 3H.**

Number of wells per genotype at each DIV, one-way ANOVA F values at each DIV, two-way ANOVA (genotype × DIV) results, and exact Tukey post hoc p values for every pairwise comparison shown in Fig. 3H (Fano factor). Two-way ANOVA (genotype × DIV): Genotype $F_{(2,227)}$ =55.78, p<0.0001; DIV $F_{(4,227)}$ =11.21, p=2.6e-8; Genotype × DIV $F_{(8,227)}$ =3.19, p=0.0019. One-way ANOVA at each DIV followed by Tukey post hoc tests; a two-way ANOVA (genotype × DIV) tested effects across days in vitro. Recording days differed by one day between litters, so adjacent DIVs were pooled; DIV13, 21 and 27 denote DIV13/14, DIV20/21 and DIV27/28 (x-axis labels 14, 20 and 28 in Fig. 3). N = independent wells (cultures). Shaded cells: p<0.05. * p<0.05, ** p<0.01, *** p<0.001.
